# Gene Conversion Facilitates the Adaptive Evolution of Self-Resistance in Highly Toxic Newts

**DOI:** 10.1101/2021.03.25.437018

**Authors:** Kerry L. Gendreau, Angela D. Hornsby, Michael T. J. Hague, Joel W. McGlothlin

## Abstract

Reconstructing the histories of complex adaptations and identifying the evolutionary mechanisms underlying their origins are two of the primary goals of evolutionary biology. *Taricha* newts, which contain high concentrations of the deadly toxin tetrodotoxin (TTX) as an antipredator defense, have evolved resistance to self-intoxication, which is a complex adaptation requiring changes in six paralogs of the voltage-gated sodium channel (Na_v_) gene family, the physiological target of TTX. Here, we reconstruct the origins of TTX self-resistance by sequencing the entire Na_v_ gene family in newts and related salamanders. We show that moderate TTX resistance evolved early in the salamander lineage in three of the six Na_v_ paralogs, preceding the proposed appearance of tetrodotoxic newts by ∼100 million years. TTX-bearing newts possess additional unique substitutions across the entire Na_v_ gene family that provide physiological TTX resistance. These substitutions coincide with signatures of positive selection and relaxed purifying selection, as well as gene conversion events, that together likely facilitated their evolution. We also identify a novel exon duplication within Na_v_1.4 encoding an expressed TTX-binding site. Two resistance-conferring changes within newts appear to have spread via nonallelic gene conversion: in one case, one codon was copied between paralogs, and in the second, multiple substitutions were homogenized between the duplicate exons of Na_v_1.4. Our results demonstrate that gene conversion can accelerate the coordinated evolution of gene families in response to a common selection pressure.

## Introduction

Fitting evolutionary models to molecular sequences in a phylogenetic context can help piece together the key steps in adaptive evolution and uncover the relative contributions of selection and other evolutionary mechanisms to adaptive phenotypic evolution (Smith et al. 2020). Comparative studies of convergence, or the repeated evolution of characters within different lineages undergoing the same environmental challenges, provide powerful evidence of both adaptation and connections between genetic and phenotypic change (Losos 2011). Investigations into the molecular basis of convergence have revealed multiple occurrences of parallelism, where different lineages have evolved changes within the same proteins, and occasionally at the same amino acid sites, in response to shared selective pressures, such as insects that have evolved the ability to feed on toxic plants (Zhen et al. 2012) and populations of ducks and humans living at high elevations (Graham and McCracken 2019). Such patterns support important roles for both positive selection and constraint in the origin of complex adaptations (reviewed by Storz 2016).

Resistance to tetrodotoxin (TTX), a potent neurotoxin, has evolved convergently in several distantly related organisms, including pufferfish, snakes, and newts (reviewed by Soong and Venkatesh 2006; Toledo et al. 2016), and therefore offers an ideal system to investigate the molecular basis of adaptive evolution (Arbuckle et al. 2017). The genetic basis of TTX resistance, which is well established in tetrodotoxic puffer fish (Jost et al. 2008) and in snakes that consume TTX-bearing prey (Geffeney et al. 2002; Feldman et al. 2012; McGlothlin et al. 2014; McGlothlin et al. 2016), involves amino acid substitutions in the toxin’s target, voltage-gated sodium channels. While voltage-gated sodium channels are commonly abbreviated as *SCNA* genes and Na_v_ proteins, hereafter we simplify this nomenclature by abbreviating both as Na_v_. Na_v_ channels are responsible for the initiation and propagation of action potentials in excitable cells and are composed of four domains (DI–DIV), each comprising six transmembrane helices and a pore-loop region (the P-loop; Fux et al. 2018). The four P-loops, one in each domain, form a pore within the membranes of excitable cells to selectively allow sodium ions to cross when the channel is open. TTX exerts its effects by binding to the P-loops of sensitive channels and preventing sodium entry into cells, thus blocking action potentials. Gene duplication events have resulted in six Na_v_ paralogs, each with tissue-specific expression, that are shared across all tetrapods (table 1), with additional lineage-specific duplications occurring in amniotes (Widmark et al. 2011; Zakon 2012). Because the structure of these paralogs is highly conserved, each has the potential to be blocked by TTX if it lacks resistance-conferring substitutions.

**Table 1.** Nomenclature for Voltage-Gated Sodium Channel Genes

| Gene name | Protein name | Tissue expression<sup>a</sup> |
| --- | --- | --- |
| **SCN1A** | Nav1.1 | Brain |
| **SCN2A** | Nav1.2 | Brain |
| **SCN3A** | Nav1.3 | Brain |
| **SCN4A** | Nav1.4 | Muscle |
| **SCN5A** | Nav1.5 | Heart |
| **SCN8A** | Nav1.6 | Brain / peripheral nervous system |

<sup>a</sup> Patterns of tissue expression are inferred from studies of gene orthologs in mammals (reviewed in Yu and Catterall 2003).

Species that possess or consume TTX must either have a full complement of resistant paralogs or otherwise shield sodium channels from contact with the toxin. Indeed, resistant substitutions are present in all eight of the Na_v_ genes within the genomes of multiple species of TTX-bearing pufferfish (from the family Tetraodontidae; Jost et al. 2008) and in six of the nine Na_v_ genes in *Thamnophis sirtalis* snakes that consume TTX-bearing *Taricha* newts (McGlothlin et al. 2014; Perry et al. 2018). The three brain channels Na_v_1.1, Na_v_1.2, and Na_v_1.3 of snakes remain TTX sensitive, but presumably are protected from TTX by the blood-brain barrier (McGlothlin et al. 2014). In snakes, the evolution of extreme TTX resistance appears to follow a predictable, stepwise substitution pattern across TTX-exposed members of the Na_v_ gene family, with substitutions in heart and peripheral nerve channels preceding those in the muscle channel gene, Na_v_1.4, which evolves resistance only in snakes locked in coevolutionary arms races with highly tetrodotoxic amphibian prey (Feldman et al. 2012; McGlothlin et al. 2016; Perry et al. 2018).

Less is known about the evolutionary history of TTX resistance in *Taricha* newts, the highly toxic coevolutionary partner of *Thamnophis* (Brodie and Brodie 1990; Brodie et al. 2002; Williams et al. 2010; Hague et al. 2020). The extreme toxicity of *Taricha*, which has been elaborated by the ongoing coevolutionary arms race with garter snakes, builds upon lower levels of toxicity that evolved ∼30 million years ago (mya) within “modern” newts (tribe Molgini; Hanifin and Gilly 2015; divergence date estimated by Hime et al. 2021). The evolution of toxicity necessitates the evolution of toxin autoresistance so that a prey species is not incapacitated by its own antipredator defense (Jost et al. 2008; Toledo et al. 2016; Tarvin et al. 2017; Márquez et al. 2019). Understanding the timing and details of this autoresistance can shed light on the genetic processes underlying the predator-prey arms race. Hanifin and Gilly (2015) compared the sequences of one sodium channel gene, the muscle paralog Na_v_1.4, across several salamander species and identified substitutions in the P-loops of DIII and DIV that provide extreme TTX resistance to the muscles of TTX-bearing newts. Importantly, the sister group of these toxic newts had substitutions in the same gene providing more moderate resistance, indicating that the evolution of autoresistance in a common ancestor paved the way for the evolution of extreme toxicity. More recently, Vaelli et al. (2020) used transcriptome sequencing to characterize the genetic basis of physiological resistance to TTX in *Taricha granulosa* and identified substitutions within TTX-binding regions in the other five Na_v_ paralogs, many of which occur in within the P-loop of DI. However, because it is unknown whether other salamander species possess TTX resistance in these paralogs, the order and timing of the evolutionary events leading to autoresistance in toxic newts is still unknown. Furthermore, no studies to date have applied evolutionary models to test for the relative importance of mechanisms such as positive selection, relaxed constraint, and interlocus gene conversion in the evolution of newt TTX resistance.

Here, we trace the evolutionary history of the entire Na_v_ gene family across the salamander phylogeny to show the order in which resistant substitutions appeared. Using published genome sequences and newly generated sequence data, we characterize the genomic structure of Na_v_ genes in newts and their relatives, inferring the timing of resistant substitutions leading to the extreme TTX resistance observed across all Na_v_ paralogs in *Taricha* newts (Vaelli et al. 2020). We estimate rates of synonymous and nonsynonymous substitution to identify positive selection. In addition, we assess the potential of nonallelic gene conversion, a process by which sequence is copied from one paralog to another (Chen et al. 2007), to act as a source of adaptive variation. Combining these data provides insight into the evolutionary mechanisms underlying the origin of a uniquely potent chemical defense.

## Results

### Genomic Structure of Voltage-Gated Sodium Channels

We used targeted sequence capture to characterize Na_v_ sequences from the genomes of five salamander species (order Urodela), including three TTX-bearing newts (family Salamandridae, subfamily Pleurodelinae, tribe Molgini), *Notophthalmus viridescens, Taricha torosa*, and *Taricha granulosa* (*n* = 3 diploid individuals of each species), and two salamanders that do not possess tetrodotoxin, *Cryptobranchus alleganiensis* (Crypotobranchidae) and *Plethodon cinereus* (Plethodontidae, *n* = 2 each). We also identified Na_v_ sequences within two publicly available salamander genome sequences: *Ambystoma mexicanum* (Ambystomatidae; Smith et al. 2019; AmexG.v6 assembly) and *Pleurodeles waltl* (Salamandridae; Elewa et al. 2017) and a full-body transcriptome from the fire salamander *Salamandra salamandra* (Salamandridae; Goedbloed et al. 2017; BioProject accession PRJNA607429), all three of which appear to lack tetrodotoxin (Hanifin 2010). The split between *Cryptobranchus* (suborder Cryptobranchoidea) and all the other salamanders in our study (members of suborder Salamandroidea) represents the most ancient division in the phylogeny of extant salamanders (∼160 mya; Hime et al. 2021).

We identified six Na_v_ genes in the genomes of all salamander species, which is consistent with observations in other amphibians (Zakon et al. 2011). Hereafter, we use the exon delineation introduced in Widmark et al. (2011), where domain I (DI) is encoded by exons 2-9, domain II (DII) is encoded by exons 13-16, domain III (DIII) is encoded by exons 18-23, and domain IV (DIV) is encoded by exons 25-26. We obtained near full-length assemblies for all paralogs (table S1); however, a few exons containing TTX-binding sites, including exon 15 (encoding the DII P-loop) of Na_v_1.2 from *N. viridescens* and exon 22 (encoding part of the DIII P-loop) of Na_v_1.2 for several newt species, were missing from our assemblies. Polymorphism was rare in our assemblies and we observed few nonsynonymous mutations within the newt genomes, but we found slightly elevated polymorphism in *N. viridescens* relative to other species (table S2). No heterozygosity or nonsynonymous polymorphisms were observed within any of the known TTX-binding P-loop regions within any of the species sequenced for this study.

Synteny of Na_v_ genes in *A. mexicanum* is conserved relative to other tetrapods (fig. S1), which allowed us to use the *A. mexicanum* sequences as a baseline to confidently identify Na_v_ paralogs in all species. Three of the paralogs, Na_v_1.1, Na_v_1.2, and Na_v_1.3 are arrayed in tandem on *A. mexicanum* chromosome 9, with Na_v_1.2 inverted relative to its neighboring paralogs (table 2, fig. S1). The additional three paralogs, Na_v_1.4, Na_v_1.5, and Na_v_1.6, are each located on separate chromosomes (table 2). In the gene family tree built from amino acid sequences, all salamander Na_v_ proteins formed a monophyletic clade (bootstrap support > 98%) with the corresponding orthologs from the genomes of the frogs *Xenopus tropicalis* and *Nanorana parkeri*, which we included as outgroups (fig. 1). The gene family tree constructed from the nucleotide coding sequences of these genes yielded a similar topology, with each salamander Na_v_ ortholog forming a monophyletic clade. However, in the nucleotide gene family tree, the three *X. tropicalis* nerve channels Na_v_1.1, Na_v_1.2, and Na_v_1.3 formed a monophyletic clade that is distinct from the salamander sequences (bootstrap support 86%; fig. S2). The same tree topology was resolved when partitioning for the third codon position (see Supplemental Data on Dryad).

**Table 2.** Locations of Na_v_ Genes in the *Ambystoma mexicanum* AmexG.v6 Genome Assembly.

| Gene | Chromosome | Start | End | Strand | Length (bp) |
| --- | --- | --- | --- | --- | --- |
| Na<sub>v</sub>1.1 | 9q | 503,107,904 | 503,797,933 | + | 690,029 |
| Na<sub>v</sub>1.2 | 9q | 507,685,503 | 508,688,108 | - | 1,002,605 |
| Na<sub>v</sub>1.3 | 9q | 509,830,827 | 510,576,927 | + | 746,100 |
| Na<sub>v</sub>1.4 | 13q | 113,343,821 | 115,707,581 | + | 2,363,760 |
| Na<sub>v</sub>1.4 exon 26b | 13q | 115,891,071 | 115,892,251 | + | 1,180 |
| Na<sub>v</sub>1.5 | 2p | 562,159,042 | 563,030,267 | - | 871,225 |
| Na<sub>v</sub>1.6 | 3q | 465,212,114 | 466,692,047 | - | 1,479,933 |

**Fig. 1.**
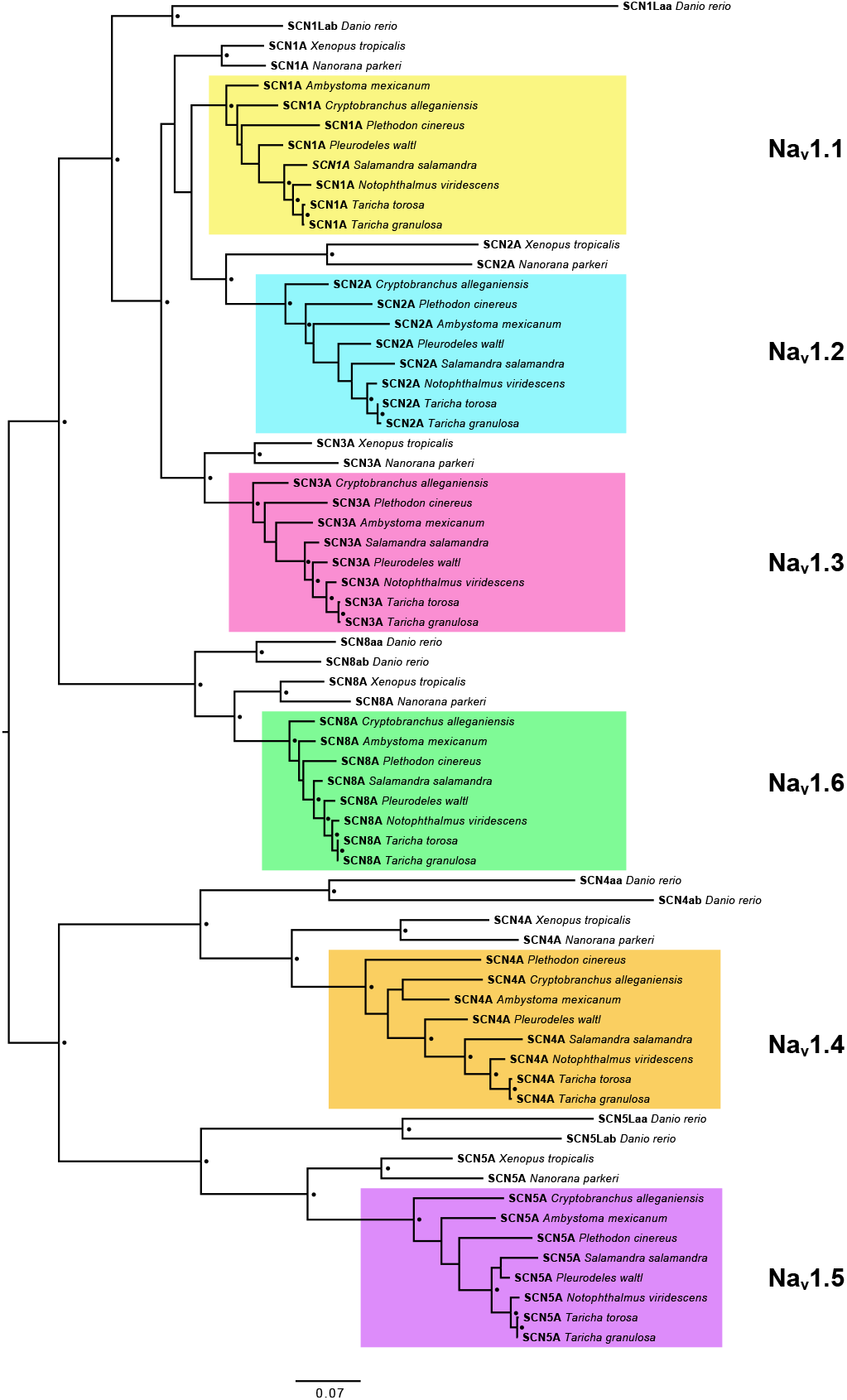
Evolutionary relationships among salamander voltage-gated sodium channels. Midpoint rooted maximum likelihood tree constructed from alignment of 2188 amino acids from full coding sequence translations of salamander sodium channel genes, including sequences from two frogs (*Nanorana parkeri* and *Xenopus tropicalis*) and one fish (*Danio rerio*) as outgroups. Black circles indicate nodes with >90% bootstrap support.

### Partial Duplication of Na_v_1.4 and Evolution of TTX Resistance in Duplicated Domains

Our search of the *A. mexicanum* genome revealed a partial tandem duplication of the 3’ end of the Na_v_1.4 gene, including the full coding region of exon 26, located ∼180,000 base pairs downstream of the full-length Na_v_1.4 gene on the same DNA strand (table 2). Both exon 26 copies are similar in length to each other and to exon 26 of other paralogs, encoding open reading frames of approximately 390 base pairs without introduced stop codons. Exon 26 is the 3’-terminal exon of the Na_v_1.4 gene and encodes the TTX-binding P-loop region of DIV. Hereafter, we refer to the duplicate exons as 26a (more proximal to exon 25) and 26b (more distal to exon 25) and duplicate P-loop regions as DIVa (more proximal to exon 25) and DIVb (more distal to exon 25). We also found this duplicated exon in Na_v_1.4 orthologs within the genomes of salamanders *P. cinereus, P. waltl, N. viridescens, T. torosa*, and *T. granulosa* and in published transcriptomes of *Tylototriton wenxianensis* and *Bolitoglossa vallecula*, but not in the transcriptome of *Hynobius retardatus* or in the genomes of *C. alleganiensis* or the frogs *X. tropicalis* or *N. parkeri*. This pattern suggests that the duplication event likely took place after the split of Cryptobranchoidea and Salamandroidea (fig. 2). Within the *S. salamandra* transcriptome, we found four unique RNA sequences transcribed from the Na_v_1.4 locus, with alternative splicing of exon 17 and alternative encoding of either exon 26a or 26b. Genome-mapped reads of multi-tissue transcriptomes of *A. mexicanum* (Bryant et al. 2017; Caballero-Pérez et al. 2018; Nowoshilow et al. 2018) indicate that these alternative transcripts have similar expression profiles across various tissues. Taken together, these observations provide evidence that the duplication of exon 26 led to the creation of functional splice variants in these salamanders.

**Fig. 2.**
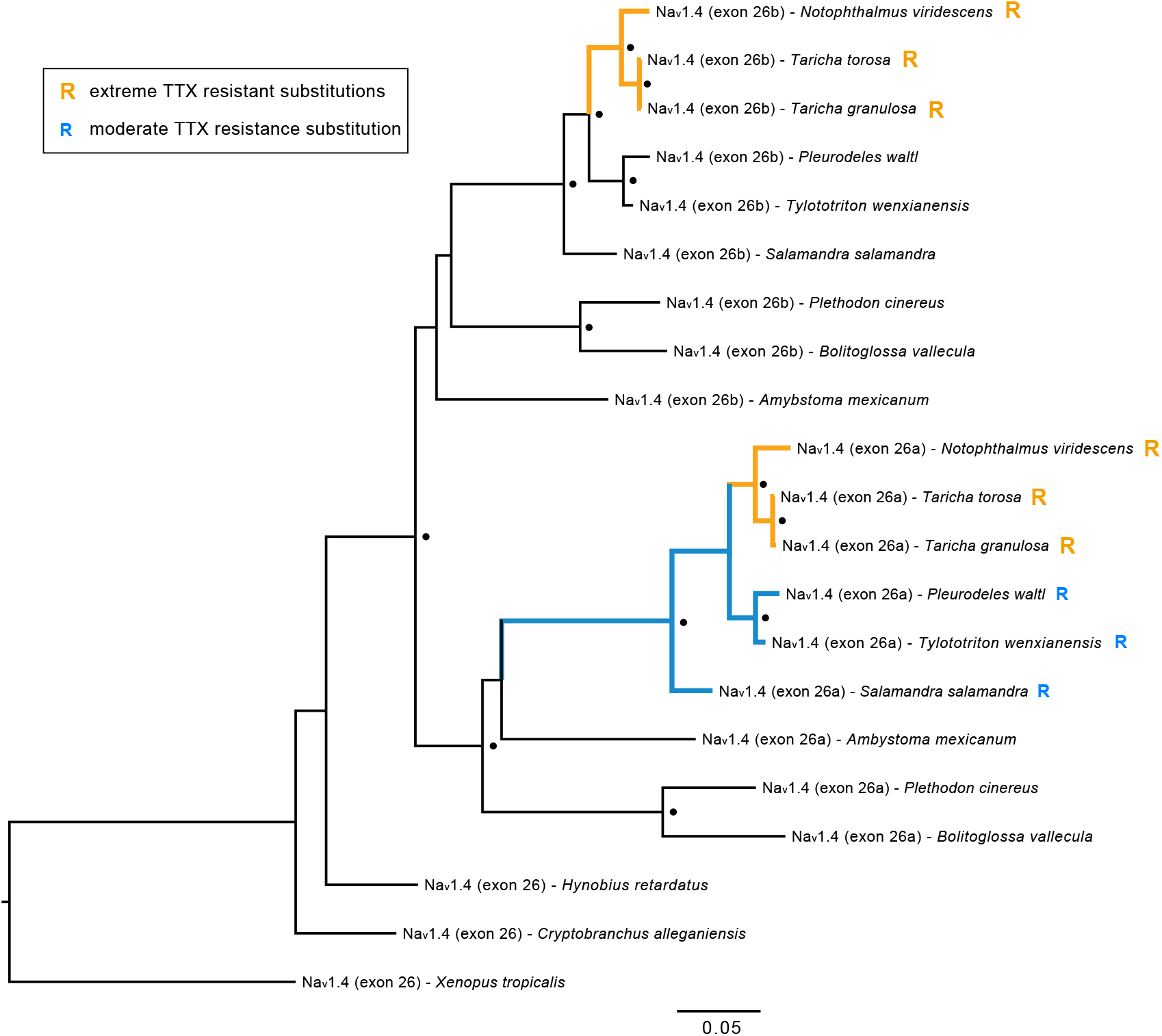
Ancestral duplication and convergent evolution of TTX resistance in Na_v_1.4 terminal exon 26. Maximum likelihood tree constructed from 1050 bp nucleotide alignment of Na_v_1.4 exon 26 identified in salamander genomes and transcriptomes. “Exon 26a” and “exon 26b” in tip labels refer to the exon copy more proximal and more distal to exon 25, respectively. Black circles indicate node bootstrap support >95%. “R” at tips indicates the presence of substitutions conferring extreme (orange) and moderate (blue) TTX resistance. The same tree topology was resolved using an alignment of amino acids translations of these sequences.

While numerous nonsynonymous substitutions differentiated the duplicated Na_v_1.4 exon 26 sequences relative to the original sequences within each genome, we found that identical substitutions conferring extreme TTX-resistance to toxic newts (Hanifin and Gilly 2015) were present in both exons from the genomes of all three TTX-bearing newts but not in other, less toxic salamander species (fig. 2). Also consistent with the results of Hanifin and Gilly (2015), we found resistant substitutions conferring moderate TTX resistance in exon 26a of *P. waltl, T. wenxianensis*, and *S. salamandra*; however, we observed no resistant substitutions in exon 26b outside of the toxic newt clade.

### Evolution of TTX Resistance in Salamanders

We characterized levels of TTX resistance in each Na_v_ paralog as extreme, moderate, and TTX-sensitive based on previous site-directed mutagenesis experiments in which substitutions were introduced to TTX-sensitive Na_v_ channels and cross-membrane Na^+^ current was measured *in vitro* in the presence and absence of TTX (table 3). Our results confirm that *T. granulosa* has six paralogs with extreme TTX resistance (table 3, fig. 3). Our findings are consistent with those reported by Vaelli et al. (2020) with one exception: we associate DIV (encoded by exon 26) substitutions A1529G, G1533V, and G1533A with Na_v_1.1 and Q1524E, G1533R, and G1533Q with Na_v_1.2 based on synteny mapping (fig. S1), gene trees (figs. 1, S2), and a phylogeny created from a coding sequence alignment of exon 26 (fig. S3), whereas the previous study reversed these assignments. We also show that substitutions with extreme TTX resistance are present in all six Na_v_ paralogs in two other species of highly toxic newt, *T. torosa* and *N. viridescens*, indicating that the common ancestor of these three species possessed extreme TTX resistance. Many of the substitutions in toxic newts parallel those found in TTX-bearing fish and in snakes that consume tetrodotoxic amphibians (table 3).

**Table 3.** List of TTX Resistance-Conferring Substitutions Observed in Salamanders

| Substitution<sup>a</sup> | Tetrodotoxic newts | Non-tetrodotoxic salamanders | Pufferfish and snakes<sup>b</sup> | Fold-change in TTX sensitivity | Resistance category | Citation |
| --- | --- | --- | --- | --- | --- | --- |
| Y401C (DI) | Na<sub>v</sub>1.1<br>Na<sub>v</sub>1.2 | Na<sub>v</sub>1.5 | Pufferfish:<br>Na<sub>v</sub>1.4a<br>Na<sub>v</sub>1.5La<br>Na<sub>v</sub>1.5Lb | ~2500× | Extreme | Venkatesh et al. (2005); Jost et al. (2008) |
| Y401A (DI) | Na<sub>v</sub>1.1<br>Na<sub>v</sub>1.3<br>Na<sub>v</sub>1.6 | Na<sub>v</sub>1.1 | Pufferfish:<br>Na<sub>v</sub>1.1La<br>Na<sub>v</sub>1.6b | >600× | Extreme | Jost et al. (2008); Vaelli et al. (2020) |
| Y401S (DI) | Na<sub>v</sub>1.5 | Na<sub>v</sub>1.5 | - | ~7000× | Extreme | Leffler et al. (2005) |
| D1532S (DIV)<br>G1533D (DIV) | Na<sub>v</sub>1.4 |  | Snake:<br>Na<sub>v</sub>1.4 | - | Extreme<sup>c</sup> | Feldman et al. (2012); Hanifin and Gilly (2015) |
| E758D (DII) | - | Na<sub>v</sub>1.5 | Pufferfish:<br>Na<sub>v</sub>1.4b | ~3000× | Extreme | Bricelj et al. (2005) |
| M1240T (DIII) | Na<sub>v</sub>1.1<br>Na<sub>v</sub>1.4 | Na<sub>v</sub>1.1<br>Na<sub>v</sub>1.5 | Pufferfish:<br>Na<sub>v</sub>1.1La<br>Na<sub>v</sub>1.1Lb<br>Na<sub>v</sub>1.4a<br>Na<sub>v</sub>1.4b | ~15× | Moderate | Jost et al. (2008) |
| V1233I (DIII) | Na<sub>v</sub>1.6 | Na<sub>v</sub>1.6 | Snake:<br>Na<sub>v</sub>1.6 | ~2× | Moderate | McGlothlin et al. (2014); Vaelli et al. (2020) |
| I1525V (DIV) | Na<sub>v</sub>1.6 | Na<sub>v</sub>1.5<br>Na<sub>v</sub>1.6 | Snake:<br>Na<sub>v</sub>1.6 | ~2× | Moderate | Geffeney et al. (2005); McGlothlin et al. (2014); Vaelli et al. (2020) |
| I1525S (DIV) | Na<sub>v</sub>1.4 | Na<sub>v</sub>1.4 |  | - | Moderate<sup>c</sup> | Hanifin and Gilly (2015) |
| I1525T (DIV) | - | Na<sub>v</sub>1.4<br>Na<sub>v</sub>1.5 |  | ~7.7× | Moderate | Du et al. (2009) |

<sup>a</sup> Amino acid sites are in reference to rat Na<sub>v</sub>1.4 (accession number AAA41682)
<sup>b</sup> Parallel substitutions in Na<sub>v</sub> channels of TTX-bearing pufferfish and in snakes that consume TTX-bearing prey
<sup>c</sup> Resistance category determined from action potentials recorded from salamander muscle fibers

**Fig. 3.**
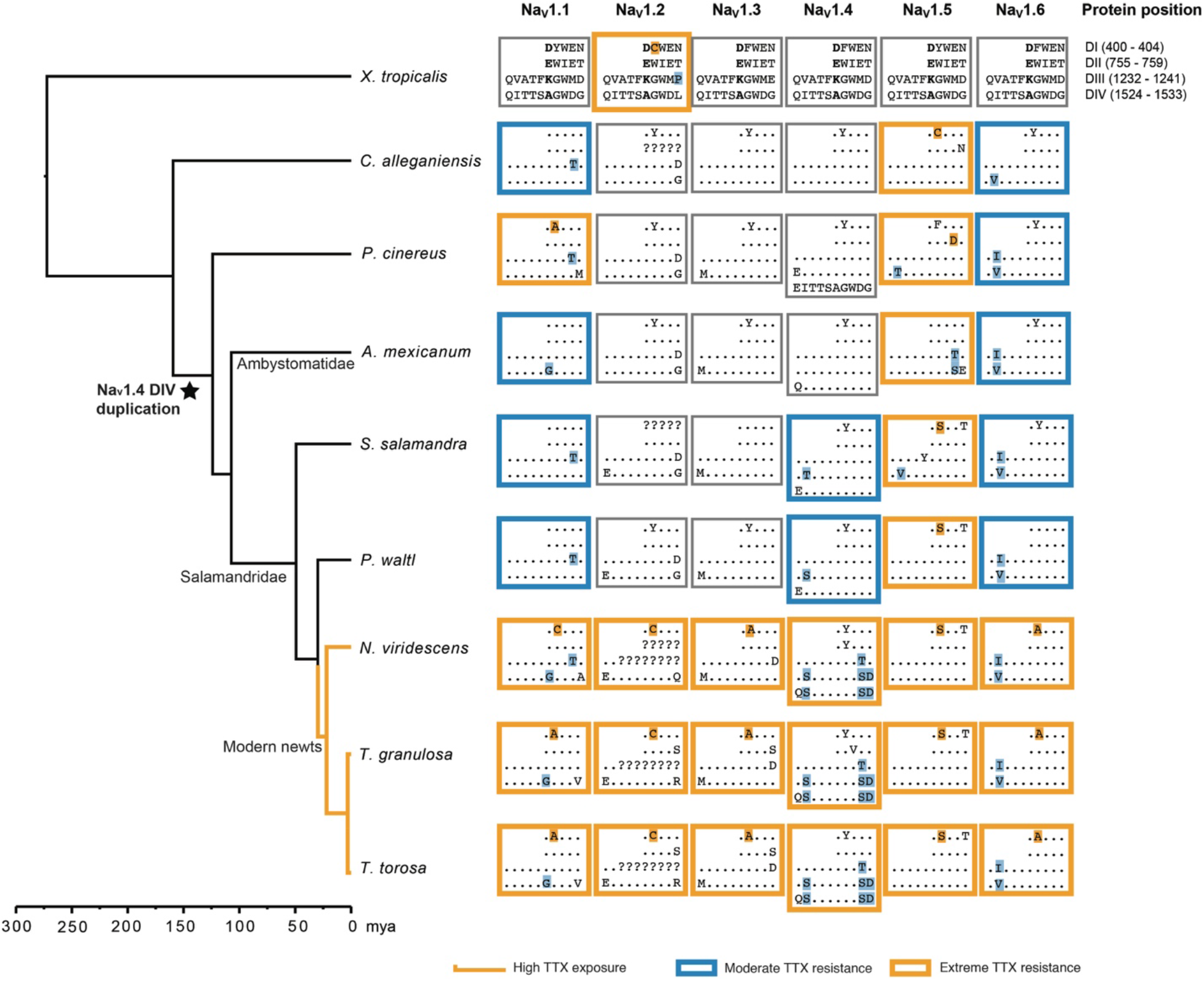
Distribution of TTX-resistance conferring substitutions in salamander voltage-gated sodium channels. Toxic newt species are indicated with orange branches (“High TTX exposure”). Amino acids associated with Na^+^ selectivity (400D, 755E, 1237K, 1529A) are shown in bold. Dots indicate identity with consensus sequences. Blue boxes indicate paralogs inferred to have moderate (∼2-15-fold) resistance, orange boxes indicate paralogs with extreme (>300-fold) resistance, and grey boxes indicate paralogs without resistance or with insufficient sequence data. Substitutions known to confer TTX resistance are highlighted with respective colors. Extreme resistance in a paralog can result from the presence of one highly resistant substitution or the combination of multiple moderately resistant substitutions. Exon duplication has led to an additional TTX-binding domain (DIVb) in Na_v_1.4 of all salamanders except *C. alleganiensis*. We did not identify the sequence encoding DIII of Na_v_1.2 for any of the newts, however Vaelli et al. (2020) report that, in *T. granulosa*, this domain is identical in amino acid sequence to other salamanders. Amino acid sites are in reference to rat Na_v_1.4 (accession number AAA41682).

No salamander species outside the clade of highly toxic newts possessed a full complement of TTX-resistant Na_v_ paralogs, indicating that the evolution of full physiological resistance coincided with the origin of extreme toxicity. However, we found at least three paralogs with moderate or extreme TTX resistance in all salamander species we examined, indicating that the evolution of TTX resistance in newts built upon more ancient changes that first appeared in their non-newt relatives. Substitutions conferring moderate or extreme resistance were observed within the heart channel Na_v_1.5 and brain/nerve channels Na_v_1.1 and Na_v_1.6 of all salamander species, with additional resistance-conferring substitutions evolving within TTX-bearing newts. As first shown by Hanifin and Gilly (2015), moderate resistance was present in the skeletal muscle channel Na_v_1.4 of *S. salamandra* and *P. waltl*, but not in the three other salamander species we examined. Although our outgroup, the frog *X. tropicalis*, also contained a highly resistant substitution in Na_v_1.2, we found no evidence for resistance in this paralog in any salamanders outside of tetrodotoxic newts.

Based on our ancestral sequence reconstructions, the most recent common ancestor of all salamanders had three TTX resistant sodium channels: Na_v_1.1 (brain, moderately resistant), Na_v_1.5 (heart, highly resistant), and Na_v_1.6 (brain/peripheral nerves, moderately resistant; figs. 3, S4-S9). Moderate resistance in the muscle channel Na_v_1.4 appeared between 75–130 mya, after the divergence of Ambystomatidae and Salamandridae (the family consisting of true salamanders and newts; Hanifin and Gilly 2015). This gain in muscle resistance coincided with the appearance of two highly resistant substitutions in DI of Na_v_1.5, which are present in all Salamandridae. Extreme TTX resistance across all Na_v_ paralogs evolved more recently, occurring approximately 30 mya, after the split between primitive newts, which include *Pleurodeles*, and modern newts (Hime et al. 2021), which include all TTX-bearing species. Over this time period, TTX resistance evolved in Na_v_1.2 and Na_v_1.3, and multiple additional resistant mutations appeared and became fixed in Na_v_1.1, Na_v_1.4, and Na_v_1.6.

### Selective Regimes and Evolutionary Rates

In order to characterize the selective regimes acting on Na_v_ genes, we used the codeml program in PAML (Yang 2007) to fit models of selection to Na_v_ codon alignments and compared nested models using likelihood ratio tests (LRTs). We tested for site-specific positive selection within all amphibians by comparing two sets of nested models: one set using a discrete distribution of ω values either with or without positive selection (M1a vs. M2a; Yang et al. 2005); and another set fitting a continuous distribution of ω values under purifying selection only, or adding categories of unconstrained evolution and positive selection (M7 vs. M8 and M8a vs. M8; Yang et al. 2000). Parameter estimates and LRT results for site models are summarized in table S3. To test for selection within toxic newts, we fit branch (Yang 1998) and branch-site models (Zhang et al. 2005) to our datasets, which allowed ω to vary both among codon sites and between toxic newts and other amphibians (summarized in table S4).

All models estimated relatively low ratios of nonsynonymous to synonymous substitution (d_N_/d_S_, or ω ratios) for all Na_v_ paralogs (average ω ratios from branch models ranged from 0.05 to 0.25), indicating pervasive purifying selection. Based on LRTs comparing one ω-value models (which allow only for one ω value across the entire phylogeny) to branch models (allowing for a different ω ratio in the toxic newt clade relative to other amphibians), we found that ω ratios were significantly (P < 0.01) higher in toxic newts for Na_v_1.1, Na_v_1.3, and Na_v_1.4 and non-significant (P > 0.01) for Na_v_1.2, Na_v_1.5, and Na_v_1.6 (fig. 4A, table S4). The largest difference in ω ratios in the branch test was observed for the muscle channel Na_v_1.4 (newt ω = 0.23; all salamanders ω = 0.10), which appears to be due to both an increase in the proportion of unconstrained sites as well as a larger number of estimated sites undergoing positive selection (fig. 4B). However, the posterior probability support for positive selection at many of these sites was low, and LRTs from branch-site models indicated significant evidence for a shift in positive selection only in paralog Na_v_1.3 (tables S3, S4).

**Fig. 4.**
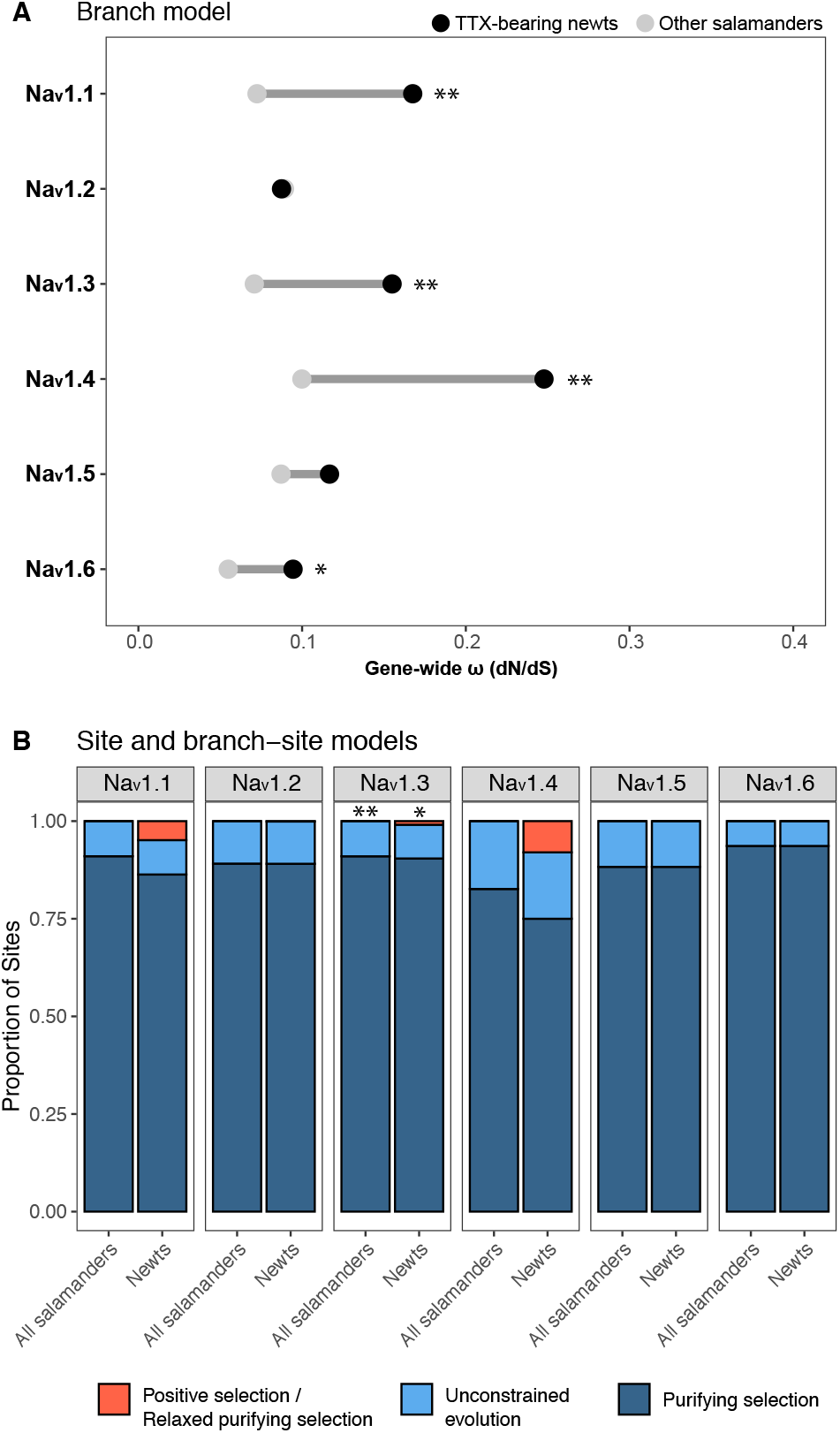
Tests for shifts in selective pressure on salamander voltage-gated sodium channels. (A) PAML branch models comparing estimates of gene-wide ω (d_N_/d_S_) within TTX-bearing newts (black circles) and other salamanders (grey circles). (B) PAML site and branch-site model estimates of proportions of sites under purifying selection (0 < ω < 0.05 in both lineages), unconstrained evolution (ω =1 in both lineages), and positive selection or relaxed purifying selection (0 < ω < 0.05 in salamanders and ω ≥ 1 in TTX-bearing newts). Significant differences based on likelihood ratio tests are indicated with * (p < 0.05) and ** (p < 0.01). Although likelihood-ratio tests were non-significant for Na_V_1.1 and Na_V_1.4 (tables S3, S4), PAML identified a large proportion of amino acid coding sites within these paralogs with elevated d_N_/d_S_ ratios in toxic newts.

Our site and branch-site models identified a number of TTX-binding sites with elevated ω ratios (tables 4, S5, S6). Because of the small number of species in our study, we had low power to detect statistically significant positive selection (Yang et al. 2000; Anisimova et al. 2001; Kosakovsky Pond and Frost 2005), and the posterior probabilities provided low-to-moderate support for positive selection at most sites. We also note that the false positive rates of site selection models can be inflated due to gene conversion events (Casola and Hahn 2009), among-site variation in d_S_ (Kosakovsky Pond and Frost 2005), and multinucleotide mutations (Venkat et al. 2018). However, results were consistent between the M2a and M8 models, and the codons that were identified suggest that positive selection may have been important for observed substitutions in TTX-binding regions.

**Table 4.** P-loop Sites with Elevated ω Values in All Salamanders and in Toxic Newts.

|  | Nav1.1 | Nav1.2 | Nav1.3 | Nav1.4 | Nav1.5 | Nav1.6 |
| --- | --- | --- | --- | --- | --- | --- |
| Tissue expression | Brain | Brain | Brain | Muscle | Heart | Brain / PNS |
| P-loop sites under positive selection in all salamanders<sup>b,d</sup> | Y401A/C (DI)<sup>a</sup><br>G1533V (DIV)<sup>a</sup> | G1533R (DIV)<sup>a</sup> | Y401A (DI)<sup>a</sup> | - | 739 (DII) | Y401A (DI)<sup>a</sup> |
| P-loop sites under positive selection in toxic newts<sup>c,d</sup> | 1224 (DIII)<br>A1529G (DIV)<sup>a</sup> | - | T759S (DII) | W756Y (DII)<br>M1240T (DIII)<sup>a</sup><br>1517 (DIV)<br>1519 (DIV)<br>D1532S (DIV)<sup>*,a</sup> | - | - |

\* Empirical Bayes posterior probability > 0.95
<sup>a</sup> Non-conservative amino acid substitutions associated with TTX resistance in toxic newts
<sup>b</sup> Sites identified with site models using empirical Bayes method (posterior probability > 0.50)
<sup>c</sup> Sites identified with branch-site models using empirical Bayes method (posterior probability > 0.50)
<sup>d</sup> Because we had low power to detect positive selection at specific sites due to the small number of species in our study, these results should be interpreted with caution

Within the brain channels Na_v_1.1, Na_v_1.3, and Na_v_1.6, putative positive selection was detected by site models at site 401 within the DI P-loop, indicating selection acting across all salamanders rather than specifically within toxic newts (table 4). Replacement of the aromatic amino acid at site 401 with a non-aromatic amino acid can substantially impact TTX binding capacity (Leffler et al. 2005; Venkatesh et al. 2005; Vaelli et al. 2020), and we observed non-aromatic substitutions at this site within all brain channels of highly toxic newts (fig. 3, table 3). The codon sequence for site 401 was variable across many salamanders lacking TTX; however, almost all of the nonsynonymous changes observed outside of TTX-bearing newts were biochemically conservative (both phenylalanine and tyrosine are aromatic and do not affect TTX binding; Sunami et al. 2000), with the exception of the 401A observed in *P. cinereus* Na_v_1.1. This conservative variation likely contributes to the signal of diversifying selection acting on this codon in less toxic salamander lineages. Site models also suggested positive selection acting on site 1533 in DIV of Na_v_1.1 and Na_v_1.2 (table 4). Although the substitutions present at site 1533 in these newt paralogs have not been tested experimentally for their effects on TTX binding, Maruta et al. (2008) showed that a G1533T substitution at this site led to a moderate (∼2-3-fold) decrease in TTX binding affinity, and substitutions at this site are common in TTX-resistant channels (Geffeney et al. 2005; Jost et al. 2008; Feldman et al. 2012; McGlothlin et al. 2016), suggesting that these non-synonymous changes likely also reduce TTX binding affinity in newts.

Within the toxic newt lineage, we identified putative positive selection acting on three known TTX-binding sites: 1240 and 1532 in DIII and DIV of muscle channel Na_v_1.4 and 1529 in DIV of brain channel Na_v_1.1 (table 4). In Na_v_1.4, sites 1240 and 1532 contain resistance-conferring substitutions exclusively within toxic newts and these substitutions have been associated with extreme TTX resistance in *Taricha* muscle fibers (Hanifin and Gilly 2015). In Na_v_1.1, site 1529 encodes part of the Na^+^ selectivity filter (comprised of interacting amino acids DEKA – sites 400, 755, 1237, and 1529), which is highly conserved across Na_v_ paralogs. An A1529G substitution (resulting in a DEKG filter) is present in Na_v_1.1 of *A. mexicanum* and all members of the toxic newt clade. While this substitution does not appear to affect Na^+^ selectivity or to be sufficient in preventing TTX from binding (Jost et al. 2008), it may alter channel firing properties, as this A1529G substitution resulted in substantially higher Na^+^ currents relative to the wild-type when introduced into a mammalian Na_v_1.4 channel (Jost et al. 2008). The same alanine to glycine substitution has been observed in Na_v_ channels of TTX-bearing flatworms (Jeziorski et al. 1997) and pufferfish (Jost et al. 2008), which suggests that it may play a role in TTX resistance in these organisms.

Outside of TTX-binding regions, we found evidence for putative positive selection in similar regions across multiple Na_v_ paralogs (tables S5, S6). For Na_v_1.4, the majority of sites under positive selection reside in terminal exon 26a that encodes the DIVa P-loop (exon 26b was excluded from this analysis due to its absence in some species). For most other paralogs, the largest clusters of sites with elevated d_N_/d_S_ ratios were within the DI L5 turret (the extracellular loop upstream of the DI P-loop, encoded by exons 6 and 7) and the DIII L5 turret upstream of the DIII P-loop encoded by exon 21.

These sites may facilitate interaction with other proteins, or alternatively, some of these sites identified by the branch-site model may be selected to compensate for biochemical changes produced by TTX-resistant mutations.

### Gene Conversion Events

We also tested for evidence of nonallelic gene conversion as a mechanism of sequence evolution contributing to adaptive evolution in Na_v_ genes using the program GENECONV. Nonallelic or ectopic gene conversion results from an interlocus exchange of DNA that can occur between closely related sequences during double-stranded break repair (Hansen et al. 2000). GENECONV uses the information in a multiple sequence alignment to identify regions of similarity shared between two sequences that is higher than expected by chance based on comparisons to permuted alignments (Sawyer 1999). We selected this program to detect gene conversion because of its low false positive rates and robustness to shifts in selective pressure (Posada and Crandall 2001; Bay and Bielawski 2011). Because nonallelic gene conversion is more likely to occur between paralogs residing on the same chromosome (Semple and Wolfe 1999; Drouin 2002; Benovoy and Drouin 2009), we limited our search to events between the tandem duplicates Na_v_1.1, Na_v_1.2, and Na_v_1.3 and between exons 26a and 26b of Na_v_1.4 within each salamander genome. While few gene conversion events were detected within each species by GENECONV, all of the regions that were detected within *Taricha* newts contain TTX-binding sites with substitutions associated with TTX resistance, including the DIVa and DIVb P-loops of Na_v_1.4 (fig. 5, table 5). We observed three TTX resistant amino acids (I1525S, D1532S, G1533D) within the DIVa and DIVb P-loops of Na_v_1.4 in toxic newt genomes. In contrast, *P. waltl*, a closely related but non-tetrodotoxic newt, contains one moderately TTX-resistant amino acid (I1525S) in the DIVa P-loop and no resistant amino acids in the DIVb P-loop (fig. 5A). These differences involve four identical nucleotide changes at homologous sites in both exon duplicates. Our short reads from the genomes of *N. viridescens* and *Taricha* newts mapped onto each of these exon assemblies across putative recombination break points with high (> 50-fold) coverage, lending support for sequence convergence rather than an assembly error. We did not detect gene conversion between these exons within the genomes of *T. granulosa* or *N. viridescens*; however, this may be due to the low power of GENECONV to detect conversion (Bay and Bielawski 2011), particularly in the presence of low sequence diversity and when the conversion tract is shorter than ∼100 bp (Posada and Crandall 2001; McGrath et al. 2009). Together, these results suggest that the three resistant amino acids accumulated together in one exon copy followed by conversion of the other exon ∼30 mya in a toxic newt ancestor.

**Fig. 5.**
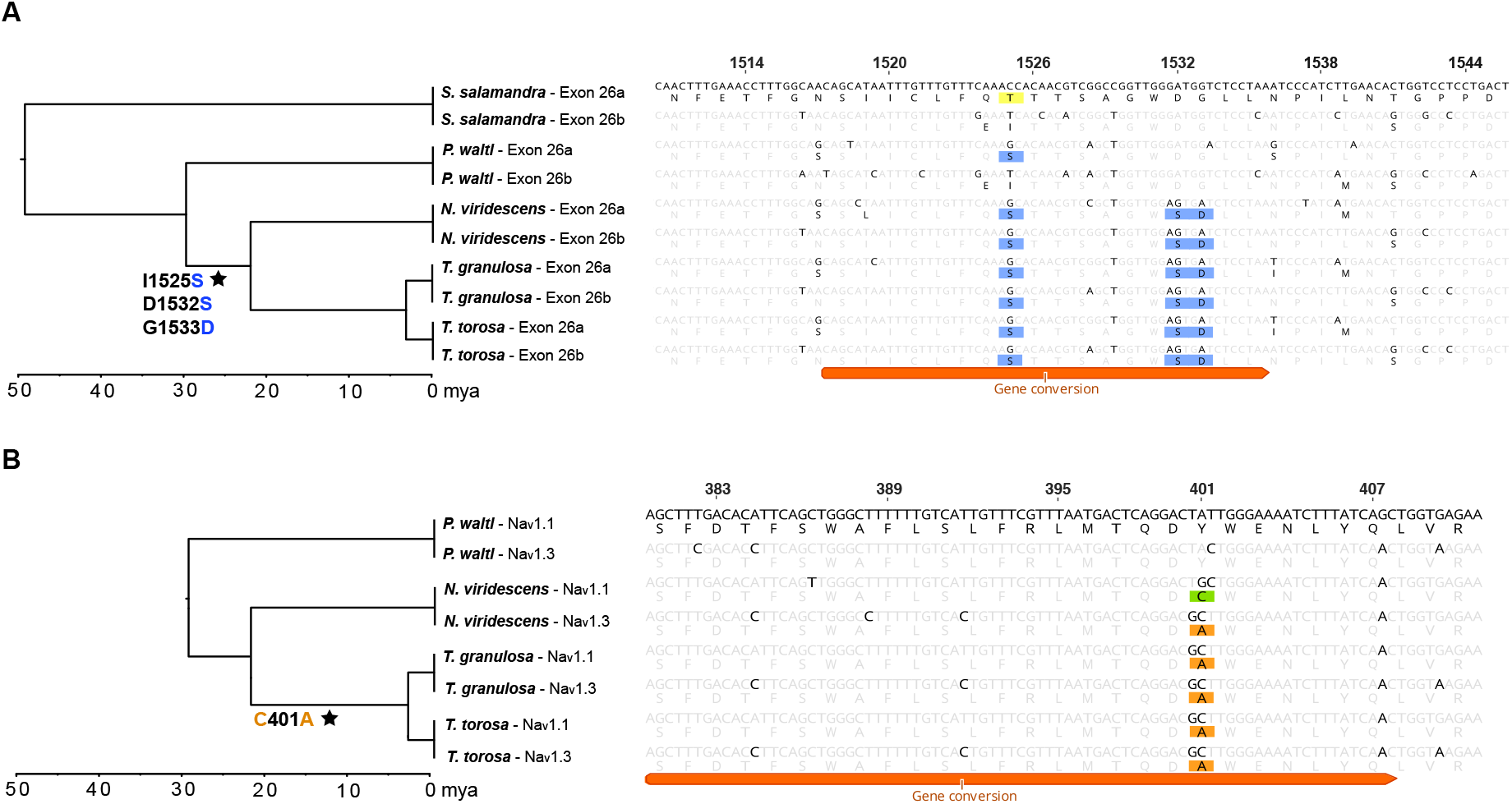
Gene conversion within TTX binding regions. (A) Gene conversion was detected between DIVa and DIVb P-loops within duplicate exons 26a and 26b of Na_v_1.4 in *T. torosa*. Substitutions conferring TTX resistance are highlighted in yellow and blue. While one moderately resistant substitution is present in DIVa of less toxic salamanders (1525T/S), three identical substitutions are present in both DIVa and DIVb of highly toxic newts. (B) Gene conversion was also detected between the DI P-loops of paralogs Na_v_1.1 and Na_v_1.3 in both *T. torosa* and *T. granulosa*. In *Notophthalmus*, both paralogs have different resistant substitutions (401C in Na_v_1.1 – highlighted in green; 401A in Na_v_1.3 – highlighted in orange). This putative gene conversion may have homogenized TTX-resistant substitutions among these paralogs within *Taricha* newts. Site numbers are in reference to amino acids of rat Na_v_1.4 (accession number AAA41682). Stars at branches indicate the inferred transfer of substitutions associated with moderate resistance (blue text) or extreme resistance (orange text) via non-allelic gene conversion.

**Table 5.** Nonallelic Gene Conversion Events

| **Nav1.1 – Nav1.2 – Nav1.3 Coding Sequences** |  |  |  |  |  |  |  |  |  |  |
| --- | --- | --- | --- | --- | --- | --- | --- | --- | --- | --- |
| **Species** | **Gene 1 – Gene 2** | **BC p-value<sup>a</sup>** | **Protein begin<sup>b</sup>** | **Protein end<sup>b</sup>** | **P-loop** | **Exon** | **Length (bp)** | **Total poly (bp)<sup>c</sup>** | **Num diffs (bp)<sup>d</sup>** | **Total diffs (bp)<sup>e</sup>** |
| *A. mexicanum* | Nav1.1 - Nav1.2 | 0.01 | 12 | 42 | - | 1 | 93 | 48 | 5 | 794 |
|  | Nav1.1 - Nav1.3 | 0.01 | 45 | 84 | - | 1 | 128 | 54 | 5 | 722 |
|  | Nav1.2 - Nav1.3 | 0.00 | 203 | 405 | DI | 5; 6; 7; 8<br>19; 20; | >626 | 74 | 2 | 902 |
|  | Nav1.1 - Nav1.2 | 0.00 | 1059 | 1193 | - | 21 | >407 | 134 | 25 | 794 |
|  | Nav1.1 - Nav1.3 | 0.02 | 1104 | 1144 | - | 20 | 127 | 37 | 1 | 722 |
|  | **Nav1.1 - Nav1.2<sup>f</sup>** | **0.02** | **1208** | **1232** | **DIII** | **21** | **84** | **35** | **2** | **794** |
| *C. alleganiensis* | Nav1.1 - Nav1.2 | 0.00 | 1111 | 1232 | DIII | 20;21 | >376 | 153 | 23 | 621 |
|  | **Nav1.2 - Nav1.3<sup>f</sup>** | **0.02** | **1478** | **1611** | **DIV** | **26** | **406** | **164** | **32** | **693** |
|  | Nav1.1 - Nav1.2 | 0.00 | 1531 | 1652 | DIV | 26 | 371 | 177 | 27 | 621 |
| *N. viridescens* | Nav1.1 - Nav1.3 | 0.00 | 500 | 512 | - | 10 | 89 | 41 | 1 | 733 |
|  | Nav1.1 - Nav1.2 | 0.01 | 1104 | 1199 | - | 16 | 292 | 109 | 20 | 796 |
| *P. cinereus* | Nav1.1 - Nav1.2 | 0.00 | 12 | 32 | - | 1 | 64 | 31 | 0 | 838 |
|  | Nav1.1 - Nav1.3 | 0.00 | 362 | 499 | DI | 8; 9; 10 | >576 | 88 | 13 | 786 |
|  | Nav1.2 - Nav1.3 | 0.02 | 573 | 599 | - | 13 | 83 | 29 | 2 | 897 |
|  | Nav1.1 - Nav1.3 | 0.00 | 610 | 667 | - | 14 | 177 | 68 | 8 | 786 |
|  | Nav1.1 - Nav1.2 | 0.00 | 1135 | 1305 | DIII | 20; 21;<br>22; 23;<br>24 | >524 | 138 | 9 | 838 |
|  | Nav1.1 - Nav1.3 | 0.03 | 1430 | 1473 | - | 26 | 134 | 46 | 5 | 786 |
| *P. waltl* | Nav1.1 - Nav1.3 | 0.03 | 1369 | 1386 | - | 25 | 56 | 30 | 0 | 738 |
| *T. granulosa* | **Nav1.1 - Nav1.3<sup>g</sup>** | **0.01** | **374** | **408** | **DI** | **8** | **105** | **41** | **2** | **725** |
| *T. torosa* | **Nav1.1 - Nav1.3<sup>g</sup>** | **0.01** | **374** | **408** | **DI** | **8** | **105** | **41** | **2** | **725** |

**Nav1.4 Exon 26**
| **Species** | **Gene** | **BC p-value<sup>a</sup>** | **Protein begin<sup>b</sup>** | **Protein end<sup>b</sup>** | **P-loop** | **Exon** | **Length (bp)** | **Total poly (bp)<sup>c</sup>** | **Num diffs (bp)<sup>d</sup>** | **Total diffs (bp)<sup>e</sup>** |
| --- | --- | --- | --- | --- | --- | --- | --- | --- | --- | --- |
| *P. cinereus* | Nav1.4 | 0.01 | 1509 | 1534 | DIV | 26a-26b | 77 | 26 | 0 | 154 |
| *T. torosa* | **Nav1.4<sup>g</sup>** | **0.02** | **1518** | **1536** | **DIV** | **26a-26b** | **56** | **22** | **1** | **178** |

<sup>a</sup> Global Bonferroni-corrected (BC) p-values from 10,000 simulations
<sup>b</sup> Protein begin and end sites are in reference to the rat Nav1.4 channel (accession number AAA41682)
<sup>c</sup> Polymorphism within the converted region across all species
<sup>d</sup> Pairwise differences between the two paralogs within the converted region
<sup>e</sup> Total pairwise differences across the entire length of the two paralogs
<sup>f</sup> Putative gene conversion events associated with the loss of TTX resistance
<sup>g</sup> Putative gene conversion events associated with the gain of TTX resistance

We also detected gene conversion between the DI P-loops of paralogs Na_v_1.1 and Na_v_1.3 within the genomes of both *Taricha* species (fig. 5B, table 5). TTX-resistant substitutions are identical in the DI P-loops of Na_v_1.1 and Na_v_1.3 in both *Taricha* species (401A, encoded by a GCT codon), while Na_v_1.1 of *N. viridescens* contains a different codon at this position (TGC, encoding 401C), consistent with gene conversion occurring at this locus in *Taricha*. Both Na_v_1.1 and Na_v_1.3 of non-tetrodotoxic *P. waltl* newts encode a TTX-sensitive tyrosine at this locus. It is unclear whether putative gene conversion event(s) occurred before or after resistance evolved in both paralogs. Gene conversion may have converted a non-resistant channel to a resistant channel in an ancestral *Taricha*, while *N. viridescens* independently acquired a Y401A substitution, or it may have homogenized substitutions within two channels that had previously evolved resistance in an ancestor of all toxic newts. In *Taricha* newts, the transition from either tyrosine or cysteine to alanine required multiple nucleotide substitutions in both paralogs, making gene conversion a likely explanation for the observed substitution patterns.

We detected additional gene conversion events that may have involved a non-resistant paralog acting as a donor to a resistant paralog, leading to the loss of TTX resistance in *A. mexicanum* and *C. alleganiensis* paralogs. A resistant substitution is present in the DIII P-loop of Na_v_1.1 in most salamanders but is absent in Na_v_1.1 of *A. mexicanum* (fig. 3), and we detected gene conversion in an adjacent region between Na_v_1.1 and the non-resistant Na_v_1.2 paralog within the *A. mexicanum* genome (table 5, fig. S10A). Similarly, *C. alleganiensis* Na_v_1.2 contains a non-resistant 1533G within DIV of all six paralogs, while Na_v_1.2 of *X. tropicalis* frogs encodes a putatively TTX-resistant 1533L, and gene conversion between paralogs Na_v_1.2 and Na_v_1.3 may have facilitated the loss of this substitution in salamanders (fig. S10B)

## Discussion

Here we show that TTX-bearing newts have evolved resistance to their own toxicity through multiple parallel changes in their Na_v_ genes and that, similar to snakes that consume TTX-containing prey (McGlothlin et al. 2016; Perry et al. 2018), some of the resistance in this taxon is ancient, first appearing in an early salamander. However, while substitutions conferring moderate TTX resistance are present in some non-toxic salamander genomes, only the TTX-bearing newts have substitutions conferring high resistance across all six of their Na_v_ paralogs, and several of these channels harbor multiple resistant substitutions in more than one domain. Many of the substitutions conferring resistance to toxic newts are also present in Na_v_ paralogs of TTX-resistant pufferfish (Jost et al. 2008) and snakes (McGlothlin et al. 2016). Similar to pufferfish, newts appear to require resistance within all of their brain/nerve channels in addition to their hearts and muscles. This feature apparently distinguishes toxic prey from their predators, whose brain channels lack resistant substitutions (McGlothlin et al. 2014). This molecular parallelism emphasizes the strong structural and functional constraints on this gene family, which appear to limit the evolution of TTX resistance to a small number of predictable pathways, leading to convergent and parallel changes across multiple taxa (Feldman et al. 2012). We show that the evolution of extreme TTX resistance is accompanied by a shift in signatures of selection in four out of six paralogs, with suggestive evidence of positive selection acting directly on TTX-binding sites. Finally, the evolution of physiological resistance appears to have been facilitated by at least two instances of non-allelic gene conversion, which acted to introduce TTX-resistance substitutions from one paralog (or exon duplicate) to another, providing a rare example of the facilitation of adaptive evolution via non-allelic gene conversion.

Our reconstruction of the history of TTX resistance in salamanders reveals the ancient origins of moderate resistance in nerve channels Na_v_1.1 and Na_v_1.6 and high resistance in the heart channel Na_v_1.5, which arose ∼160 mya. Resistant substitutions in the muscle channel Na_v_1.4 evolved later, becoming fixed in the clade including all newts and *S. salamandra* between 75–130 mya. This was followed by the accumulation of additional substitutions in DIII, DIVa, and DIVb of Na_v_1.4 within members of the highly toxic newt clade, providing their muscles with resistance to much higher concentrations of TTX relative to salamanders lacking these mutations (Hanifin and Gilly 2015). Substitutions in DI conferring extreme TTX resistance to the brain/nerve channels Na_v_1.1, Na_v_1.2, Na_v_1.3, and Na_v_1.6 also evolved more recently (∼30 mya) and they are limited to the toxic newt clade with only one exception, Na_v_1.1 of *P. cinereus*, which may have arisen independently in this lineage. Toxic newts also have unique substitutions in DIV of Na_v_1.1 and Na_v_1.2, which may provide additional resistance to their brain channels or may compensate for structural or functional changes resulting from resistant substitutions in DI or in other regions of the protein. The widespread presence of TTX in modern newts suggests that possession of TTX evolved in the common ancestor of this clade (Hanifin 2010). This is supported by our observation of shared TTX-resistance substitutions across all Na_v_ paralogs in highly toxic newts. However, the highly toxic newts included in this study are limited to North American species and do not include Asian or European newts (e.g. *Cynops* and *Triturus* spp.) known to have TTX (Hanifin 2010). Sequencing the Na_v_ genes of these species will reveal if extreme resistance in the nerve channels is specific to North American newts or arose earlier in an ancestral newt species.

Whether the TTX-resistance substitutions observed in newt relatives are adaptive and what selective pressures act to retain them remains to be determined. In ancestral salamandrids, resistance in three or four of the six Na_v_ proteins may have provided tolerance to low levels of TTX, facilitating the evolution of extreme toxicity in the modern newts, although it is unclear whether these substitutions arose in response to TTX exposure or as a side effect of selection for another aspect of channel function. Environmental exposure to TTX could potentially occur from TTX-bearing prey, such as terrestrial *Bipalium* flatworms (Stokes et al. 2014) or from TTX-producing bacteria (Vaelli et al. 2020). In a feeding study on *B. adventitium* flatworms, Ducey et al. (1999) demonstrated that while all salamanders rejected the flatworms when they were first presented, some habituated *Ambystoma* and *Plethodon* individuals were able to consume them with only minor symptoms of TTX poisoning, including apparent mucus production and numbing of the mouth. Although the amount of TTX present in the worms was not measured, this observation supports the conjecture that these substitutions play a protective role against consuming toxic prey.

Our power to detect sites undergoing positive selection was low due to the small number of species available for our analyses and the high sequence conservation among orthologs (Yang et al. 2000; Anisimova et al. 2001). However, our analyses provided evidence consistent with positive selection acting on TTX-binding sites, including sites implicated in the extreme TTX resistance of salamander muscle fibers. Our branch models indicate that several Na_v_ paralogs have higher ω ratios within the tetrodotoxic newt clade relative to other salamanders. This increase in ω ratio appears to be mainly the result of relaxed purifying selection rather than positive selection, as few sites were estimated to have ω > 1 by branch-site models. Muscle channel Na_v_1.4 had the highest ω ratio in toxic newts, coinciding with a relatively high proportion (∼0.06) of sites detected as undergoing positive selection in toxic newts and purifying selection in other salamanders, although the likelihood ratio test was non-significant (table S4). This pattern may derive from ongoing positive selection on Na_v_1.4 resulting from the coevolutionary arms race between newts and snakes. Increased ω ratios in toxic newts coincide with the appearance of highly TTX-resistant substitutions within the DI P-loops of Na_v_1.1, Na_v_1.3, and Na_v_1.6, DIV of Na_v_1.1, and the DIII, DIVa, and DIVb P-loops in Na_v_1.4. While the pattern of these substitutions suggests that they are adaptive changes that occurred specifically within the tetrodotoxic newt clade, our branch-site models detected positive selection on TTX-binding sites only within the DIV P-loop of Na_v_1.1 and the DIII and DIVa P-loops of Na_v_1.4, while our site models detected positive selection on site 401 within paralogs Na_v_1.1, Na_v_1.3, and Na_v_1.6 of all salamanders. This may be due to the statistically conservative nature of the branch-site model and the tendency of PAML to detect ongoing diversifying selection rather than rare positive selection events (Yang and dos Reis 2010). Furthermore, the site and branch-site models do not distinguish between biochemically conservative and non-conservative amino acid changes. Nevertheless, the detection of relatively high ω ratios within the toxic newt clade along with site-specific positive selection at known TTX-binding sites provides strong evidence that these substitutions are adaptive.

In addition to the six Na_v_ paralogs described in amphibians (Zakon 2012), our sequence data revealed the presence of a partial duplication of the Na_v_1.4 gene in salamanders that includes the entirety of exon 26, encoding the DIV P-loop, which likely occurred in an ancestor of Salamandroidea. The maintained open reading frame and shared expression patterns of transcripts encoding exons 26a and 26b in *A. mexicanum* tissues suggests that this duplicate region is functional in salamander muscles. In some insects that feed on toxic plants, the appearance of resistance-conferring substitutions is accompanied by one or more duplications of the genes that the toxin targets (Zhen et al. 2012; Petschenka et al. 2017). Petschenka et al. (2017) show that in at least one species, such gene duplication precedes resistance, and they suggest that gene duplication may help to alleviate the potential decrease in fitness incurred by the insects due to the negative pleiotropic effects of toxin-resistant mutations. Similarly, resistance in DIV of salamander Na_v_1.4 appears after the duplication of this domain, and only the TTX-bearing newts have resistant substitutions in both exon copies, lending support to this hypothesis and raising the possibility that the evolution of physiological resistance in salamanders may have been mediated in part by this genomic novelty.

We observed a rare case of the generation of adaptive variants via non-allelic gene conversion, which occurred both between the duplicated exons of Na_v_1.4 and between paralogs Na_v_1.1 and Na_v_1.3. Gene conversion is often thought to play a role in constraint, preserving the core functions of gene families (Chapman et al. 2006) and reducing deleterious mutation loads (Ohta 1989; Khakhlova and Bock 2006), and has also been associated with the diversification of major histocompatibility complex genes in mammals (Kuhner et al. 1991; Go et al. 2003) and the introduction of deleterious nonsynonymous mutations into different parts of the genome (Galtier et al. 2009; Casola et al. 2012). However, the potential for gene conversion to facilitate adaptation is less widely appreciated. Theory suggests that gene duplication and subsequent gene conversion may allow for movement between adaptive peaks via the accumulation of beneficial mutations in one gene copy that can be transferred to a favorable genetic background (Hansen et al. 2000). This process has been implicated in the adaptive evolution of hypoxia tolerance in high-altitude Tibetan wolves (Signore et al. 2019) and in heavy metal tolerance in *Arabidopsis* (Hanikenne et al. 2013). We observed a rare case of the generation of adaptive variants via non-allelic gene conversion, which occurred both between the duplicated exons of Na_v_1.4 and between paralogs Na_v_1.1 and Na_v_1.3. In the brain channel genes Na_v_1.1 and Na_v_1.3, this likely occurred between two genes that had previously evolved resistance, homogenizing the substitutions between the two paralogs in *Taricha* newts. In the muscle channel gene Na_v_1.4, three identical amino acids were present within both duplicated DIV TTX-binding domains in toxic newts, while only one resistant amino acid was present in a single DIV domain in their non-tetrodotoxic salamander relatives, suggesting that resistant substitutions accumulated in one exon copy in a toxic newt ancestor and subsequently spread to the other exon copy via gene conversion. Both Hanifin and Gilly (2015) and Du et al. (2009) have shown that the single resistant amino acid observed in less toxic salamanders confers only low levels of TTX resistance, while the combination of the three amino acids found in highly toxic newts confers extreme resistance. As splice variants encoding the alternative exons appear to share similar expression patterns in salamanders, both exon copies should require extreme resistance in species exposed to high TTX concentrations. The concerted evolution of TTX resistance among homologous P-loop domains of Na_v_1.4 may have expedited the evolution of extreme resistance in newt muscles, requiring new resistant mutations to appear in only one of these domains before being transferred to the other copy.

The degree of parallel molecular evolution among members of the Na_v_ gene family and across lineages provides insight into the constraints on Na_v_ nucleotide sequence as well as the evolvability of the TTX-resistance phenotype. Our results reveal that similar to their coevolutionary partners, *Thamnophis* garter snakes, *Taricha* newts evolved extreme TTX resistance through a stepwise process that built upon ancient changes that were in place millions of years before the arms race began. However, the pattern of TTX resistance evolution in newts also displays important differences from that of their predators. First, perhaps because of the constitutive presence of TTX, newts display extreme levels of resistance even in channels that are expressed in the central nervous system, which are protected by the blood-brain barrier in species that encounter TTX in their diet. Second, our analysis indicates that many substitutions may have become fixed relatively close to one another in evolutionary time within the clade of modern newts. This is in contrast to snakes, where key changes were separated by millions of years. Due to our lack of sampling of newt species outside of North America, however, further work is necessary to understand the timing of these changes on a finer scale. We also show that while positive selection appears to be a strong driving force of the evolution of TTX auto-resistance in newts, gene conversion may have sped up the process of adaptive evolution in some Na_v_ paralogs, and constraints have limited the possible locations and types of resistant substitutions to a small subset of realized genetic changes. Taken together, our results emphasize the interplay among selection, constraint, and historical contingency in the evolution of complex adaptations.

## Methods

### Sequencing and Annotation of Voltage-Gated Sodium Channel Paralogs

We identified Na_v_ genes in the two publicly available salamander genome assemblies, *A. mexicanum* (Smith et al. 2019; AmexG.v6 assembly) and *P. waltl* (Elewa et al. 2017), and in one full-body transcriptome from the fire salamander *S. salamandra* (Goedbloed et al. 2017; BioProject accession PRJNA607429) using the reciprocal best BLAST hit method (Moreno-Hagelsieb and Latimer 2007) with queries of Na_v_ sequences from *X. tropicalis* (Hellsten et al. 2010) and salamanders (Hanifin and Gilly 2015). We confirmed assignments of each amphibian Na_v_ paralog based on nucleotide alignments of the coding sequences with *Xenopus* sequences, as well as synteny of the chromosomal segments containing Na_v_ genes (fig. S1). These Na_v_ annotations were then used to design targeted sequencing probes and to subsequently assign paralog identity to our de novo salamander assemblies. We used Geneious v10.2.3 for sequence visualization and to create DNA and protein alignments (Kearse et al. 2012). We also performed BLASTn searches of published transcriptome assemblies of *Tylototriton wenxianensis* (PRJNA323392), *Bolitoglossa vallecula* (PRJNA419601), and *Hynobius retardatus* (Matsunami et al. 2015; PRJDB2409). However, we were unable to identify the full set of Na_v_ paralogs within these three assemblies, possibly due to their tissue specificity. Therefore, we used the sequences for probe design and to identify exon duplications in Na_v_1.4 but excluded them from PAML analyses.

In order to design targeted sequencing probes, we compiled partial and complete amphibian Na_v_ sequences obtained from NCBI databases and additional published sources (table S7) into a single FASTA file. Each individual FASTA entry included a single exon and, when available, up to 200 bp of intron sequence upstream and downstream of each exon to aid in paralog assignment. Using RepeatMasker (Smit et al. 2013-2015), we replaced transposable elements and other sequence repeats with Ns, and subsequently filtered out sequences < 120 bp in length, as well as those with more than 25% missing or ambiguous characters. We submitted this masked and filtered file, which was 465 kb in total length, to Agilent Technologies (Santa Clara, CA, USA) for custom probe design using the SureSelect tool, resulting in 7518 unique 120-mer probes. Probe sequences are available in our Dryad repository.

We obtained DNA samples from adult individuals of three TTX-bearing species (n=3 for each species): *T. torosa* from Hopland, CA, *T. granulosa* from Benton, OR, and *N. viridescens* from Mountain Lake, VA, and from two additional salamander species presumed to lack TTX (*n* = 2 for each species): *P. cinereus* collected in Mountain Lake, VA and *C. alleganiensis* collected in southwestern VA. We extracted genomic DNA using the DNeasy Blood & Tissue kit (Qiagen Inc., Valencia, CA) and prepared sequencing libraries using the SureSelect^XT^ Target Enrichment system for Illumina paired-end multiplexed sequencing from Agilent Technologies (Santa Clara, CA, USA), following the protocol for low input (200 ng) DNA samples. We used a Covaris M220 Focused-ultrasonicator to shear 200 ng of purified whole genomic DNA from each sample into ∼250 bp fragments using the following settings: duty factor 10%, peak incident power 75 w, 200 cycles per burst, and treatment time of 160 seconds. We followed the Agilent SureSelect^XT^ Target Enrichment kit protocol for end repair, adaptor ligation, amplification, hybridization and bead capture (using the custom SureSelect probes described above), indexing, and purification. We quantified the resulting enriched, indexed libraries with qPCR and combined them in equimolar concentrations into one final library pool, which was submitted to the Genomics Sequencing Center at Virginia Tech for sequencing on an Illumina MiSeq 300-cycle v2 with 150 bp paired-end reads. Prior to alignment, we trimmed Illumina reads of TruSeq3 adapter sequences, removed bases with a phred64 quality score less than 3, and filtered out subsequent reads shorter than 100 bp using Trimmomatic version 0.33 (Bolger et al. 2014). We used SPAdes 3.6.0 (Bankevich et al. 2012) to create de novo assemblies of the trimmed and filtered reads. Each Na_v_ paralog had > 10-fold sequence coverage, with an average of 32-fold coverage across all species and paralogs (table S1). After filtering, all reads had Phred Q-scores >30 for >99% of sites.

Because we designed the sequencing probes to capture only small portions of the Na_v_ intronic regions flanking exons (regions more likely to be conserved across species), each exon was assembled into a separate scaffold. For each individual from our sequencing trial, we created BLAST databases from the assembled de novo scaffolds and performed BLAST searches using single exons from the *Ambystoma* Na_v_ genes as queries. We created 26 separate nucleotide alignments, one for each individual Na_v_ exon, including sequences from *Ambystoma, Pleurodeles*, and our de novo assemblies using MAFFT v 7.450 (Katoh and Standley 2013) and created consensus neighbor-joining trees with a Tamura-Nei genetic distance model using the Geneious Tree Builder (Kearse et al. 2012) with 1000 bootstrap replicates and an 80% support threshold. The resulting tree topologies were used to assign paralog identity to each of the exons. When necessary, we included exon and intron sequences from additional species in the alignments to resolve the topology of the trees.

We concatenated all exons from each paralog into full coding sequences. Based on alignment with full-length *Xenopus* sequences, all salamander Na_v_ coding sequences collected for this study were >90% complete with the exception of two sequences from the *Salamandra salamandra* transcriptome (paralogs Na_v_1.1 and Na_v_1.2, which were 68.8% and 70.7% complete respectively; table S1). For each species, we used Na_v_ paralogs sequenced from the genome of a single individual for our downstream analyses, based on completeness of the assembly.

### Determination of TTX Resistance Levels

TTX sensitivity is commonly measured *in vitro* by using site-directed mutagenesis to introduce mutations of interest into a TTX-sensitive Na_v_ channel, followed by expression in *Xenopus* oocyte or HEK 293 cells and the application of patch-clamp whole-cell recordings to measure channel current in the presence of TTX. The fold-change in TTX sensitivity is then calculated by taking the ratio of the IC_50_ values, or the TTX concentration at which 50% of the Na_v_ channels are blocked, of mutated and wild-type channels (see table 3 for references).

Another line of evidence for resistance in salamander muscle channels (Na_v_1.4) comes from Hanifin and Gilly (2015), who estimated TTX sensitivity by recording action potentials generated from salamander muscle fibers and estimating the amount of TTX required to diminish the rise of the action potential, associating these relative changes with the presence and absence of substitutions in TTX-binding sites. They associated moderate TTX resistance (reduced sensitivity to 0.010 μM TTX) with the presence of DIII substitution M1240T and extreme resistance (low sensitivity to 300 μM TTX) with the presence of DIII and DIV substitutions M1240T, D1532S, and G1533D. We categorize levels of TTX resistance conferred by Na_v_ substitutions as extreme or moderate based on the results of Hanifin and Gilly (2015) as well as the data summarized in table 3.

### Phylogenetic Analyses and Identification of Site-Specific Evolutionary Rates

We constructed phylogenetic trees for the entire Na_v_ gene family using our *de novo* assembled sequences as well as sequences from the genomes of *A. mexicanum, P. waltl*, the whole-body transcriptome of *S. salamandra*, two frog genomes: *X. tropicalis* (Hellsten et al. 2010) and *Nanorana parkeri* (Sun et al. 2015), and one fish genome: *Danio rerio* (Howe et al. 2013). We created an amino acid alignment of translated coding sequences using MAFFT v 7.450 (Katoh and Standley 2013) and constructed maximum likelihood trees from these alignments with RAxML v8.2.11 (Stamatakis 2014) in Geneious using a GAMMA BLOSSUM62 protein model and estimated clade support with 100 bootstrap replicates. In order to improve the accuracy of the nucleotide alignment, we used this amino acid alignments to guide codon alignments with PAL2NAL v14 (Suyama et al. 2006). We identified the best fitting substitution models for the nucleotide alignment using jModelTest 2.1.10 v20160303 (Darriba et al. 2012) and constructed maximum likelihood trees of the coding sequences with RAxML v8.2.11. The two models with the lowest AIC scores were (1) the transition model with unequal base frequencies, a gamma shape parameter, and some proportion of invariable sites (TIM2+I+G) and (2) a general time-reversible model with a gamma shape parameter and a proportion of invariable sites (GTR+I+G), which is nearly identical to TIM2+I+G, but includes two additional rate parameters (Posada 2008). Because the two models are nearly equivalent and both fit our data equally well, we chose the GTR+I+G model for its ease of implementation across different programs. We repeated these methods for each individual Na_v_ paralog for downstream analyses in PAML, excluding the sequences from the outgroup species *N. parkeri* and *D. rerio*. We estimated chronograms in Figs. 3 and 5 using the chronos() function of the ape v5.4-1 package in R using a phylogenetic tree from Pyron (2014) and divergence dates from Hime et al. (2021; see Dryad repository for R code).

We estimated synonymous (d_S_) and nonsynonymous (d_N_) substitution rates, as well as the d_N_/d_S_ ratios (ω) for each paralog using codeml in PAML v4.8 (Yang 2007). In order to test for changes in selective regimes in the Na_v_ genes between salamanders and highly toxic newts (*Notophthalmus* and *Taricha* species), we fit the following models to our Na_v_ alignments: (1) one-ratio models (allowing for a single ω ratio among all sites and all branches of the phylogeny), (2) branch models (allowing for separate ω ratios for the foreground [toxic newts] and background [other salamanders]), (3) branch-site neutral models (allowing ω to vary both between toxic newts and other salamanders and among sites, with two possible categories: 0 < ω < 1 and ω = 1), and (4) branch-site models (allowing ω to vary both between toxic newts and other salamanders and among sites, with three possible categories: 0 < ω < 1, ω = 1, and ω >1, the latter being allowed only within toxic newts). To test for site-specific positive selection among all salamanders, we fit two sets of nested models: (1) discrete ω ratio models M1a vs. M2a, which allowing for ω to vary among sites, with either two possible categories: 0 < ω < 1 and ω = 1 or three possible categories: 0 < ω < 1, ω = 1, and ω > 1; and (2) continuous ω ratio models M7, M8a, and M8, which fit ω ratios of sites into a beta distribution, with ncatG (the number of ω values in the beta distribution) set to 5. We used F3×4 codon models, which estimate individual nucleotide frequencies for each of the three codon positions, and allowed codeml to estimate ω and transition-transversion rates (κ). The outputs of these models were used to estimate and compare gene-wide ω between toxic newts and salamanders (one-ratio and branch models), to identify sites under positive selection in all salamanders (site and neutral site models), and to identify sites under positive selection or with elevated ω in toxic newts relative to other salamanders (branch-site neutral and branch-site models). We also created ancestral sequence reconstructions by specifying RateAncestor = 1 within codeml configuration files. We report posterior probability estimates for ancestral sequence reconstructions using models with the highest likelihood scores (Table S3), which were the neutral site models (M8a) for all Na_v_ paralogs except Na_v_1.3, for which the site selection model M8 had the most support (Fig. S4-S9). We performed all of the above analyses both including and excluding the two *S. salamandra* paralogs with a large number of gaps (Na_v_1.1 and Na_v_1.2). Maximum likelihood parameter estimates were largely congruent using these different datasets. Therefore, we present results from the gene trees including *S. salamandra* here.

### Detection of Gene Conversion

We used the program GENECONV to detect potential nonallelic gene conversion events between Na_v_ paralogs. We used a full codon alignment of all six Na_v_ paralogs from 7 of the 8 salamanders included in the study (excluding *S. salamandra* due to missing data), but targeted our search to include only genes Na_v_1.1, Na_v_1.2, and Na_v_1.3 within each species under the assumption that gene conversion is more likely to occur between closely related sequences that reside on the same chromosome. This increased the power of detecting gene conversion from multiple pairwise comparisons. We also performed a separate search for gene conversion events between duplicate exons 26a and 26b of Na_v_1.4. For this analysis, we used a codon alignment of Na_v_1.4 exons 26a and 26b from 10 salamander species. We assigned a mismatch penalty using gscale=1 and used corrected *p*-values from 10,000 permutations to determine significance.

## Acknowledgments

We thank Miguel Vences for helpful feedback on the manuscript and for providing an early draft of the *Salamandra salamandra* transcriptome. We also thank Vincent Farallo, Edmund Brodie III, William Hopkins, and Brian Case for providing DNA samples for this study, Kaitlyn Malewicz, Mercedes Collins, and Emily Orr for assistance with lab work, John Abramyan, Edmund Brodie III, Charles Hanifin, and Andrew Kern for helpful conversations and Matthew Hahn for advice on analyses and comments on the manuscript. Hellbender (*Cryptobranchus*) samples were collected by Brian Case and William Hopkins in cooperation with the Virginia Department of Wildlife Resources (DWR) and the U.S. Forest Service, and with assistance from J.D. Kleopfer. Hellbenders were handled under scientific collecting permit #060465 and approved Virginia Tech IACUC procedures (protocol #16-162). We thank the Departments of Fish and Wildlife in Oregon and California and the Virginia DWR for scientific collecting permits to MTJH (063-18, SC-11937 and 048104 respectively). This work was supported by awards from the National Science Foundation to JWM (DEB-1457463 and IOS-1755055), and to MTJH (DEB-1601296), by an R.C. Lewontin award from the Society for the Study of Evolution to KLG, and by a graduate fellowship award from the Interfaces of Global Change program at Virginia Tech to KLG.

## Data Availability

Newly generated sequences have been submitted to the SRA database in NCBI under BioProject accession PRJNA732671. Sequence alignments and configuration files for GENECONV analyses have been deposited at https://github.com/kerrygendreau/Newt-TTX-Resistance.git and archived in a Dryad repository at http://doi.org/10.5061/dryad.kprr4xh4v.

## Notes

https://github.com/kerrygendreau/Newt-TTX-Resistance.git

### Competing Interest Statement

The authors have declared no competing interest.

