## Supplemental Figures and Tables for "Gene Conversion Facilitates the Adaptive Evolution of Self-Resistance in Highly Toxic Newts": Gendreau MBE Supporting Information 25May2021.pdf

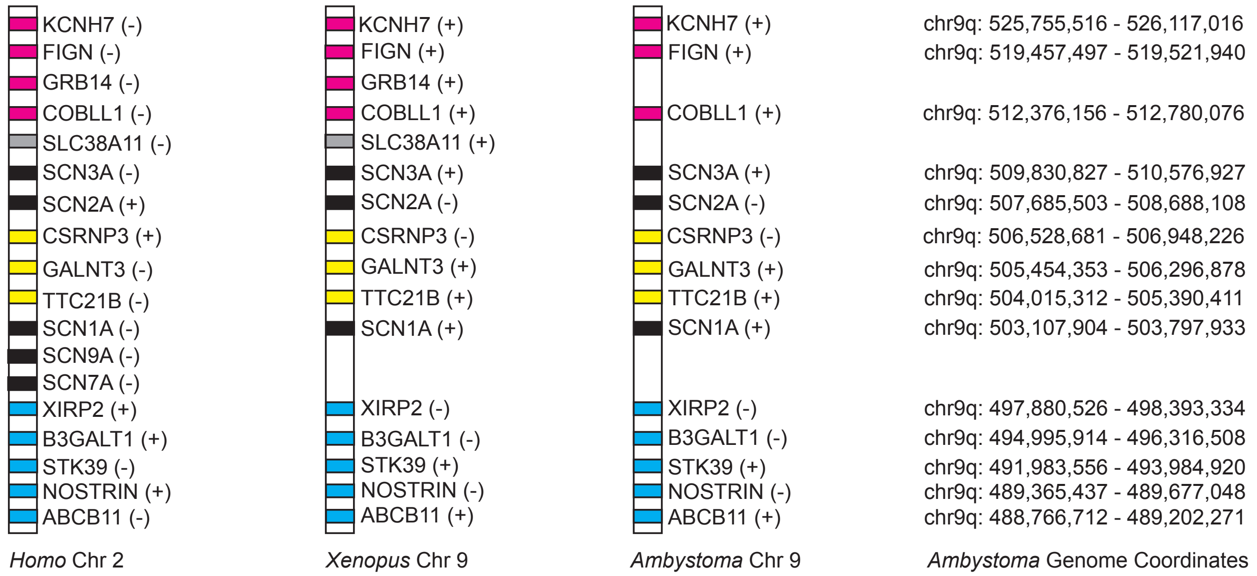


**Fig. S1.** Conserved synteny of voltage-gated sodium channel paralogs across tetrapods**.** The genetic configuration of brain/nerve voltage-gated sodium channels (*SCNA* genes) is highly conserved across three tetrapod species: humans (*Homo*), frogs (*Xenopus*), and salamanders (*Ambystoma*). Genome coordinates are based on the AmexG.v6 assembly. Symbols (+) and (-) refer to gene orientation within this reference genome.

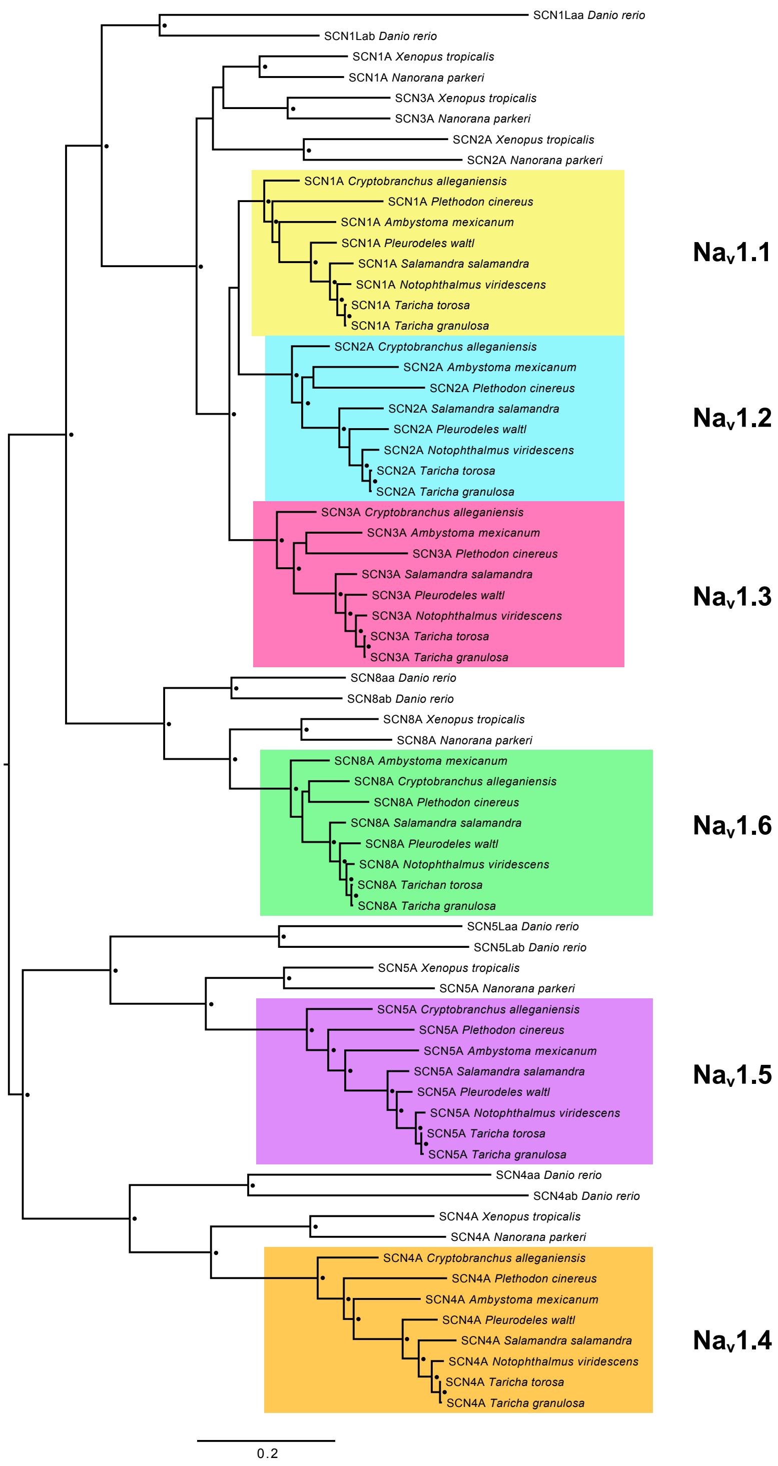


**Fig. S2.** Maximum likelihood tree constructed from 6789 bp coding sequence alignment of salamander $Na_v$ genes with coding sequences from frogs (*Nanorana parkeri* and *Xenopus tropicalis*) and fish (*Danio rerio*) as outgroups. Black circles indicate nodes with bootstrap support >90%.

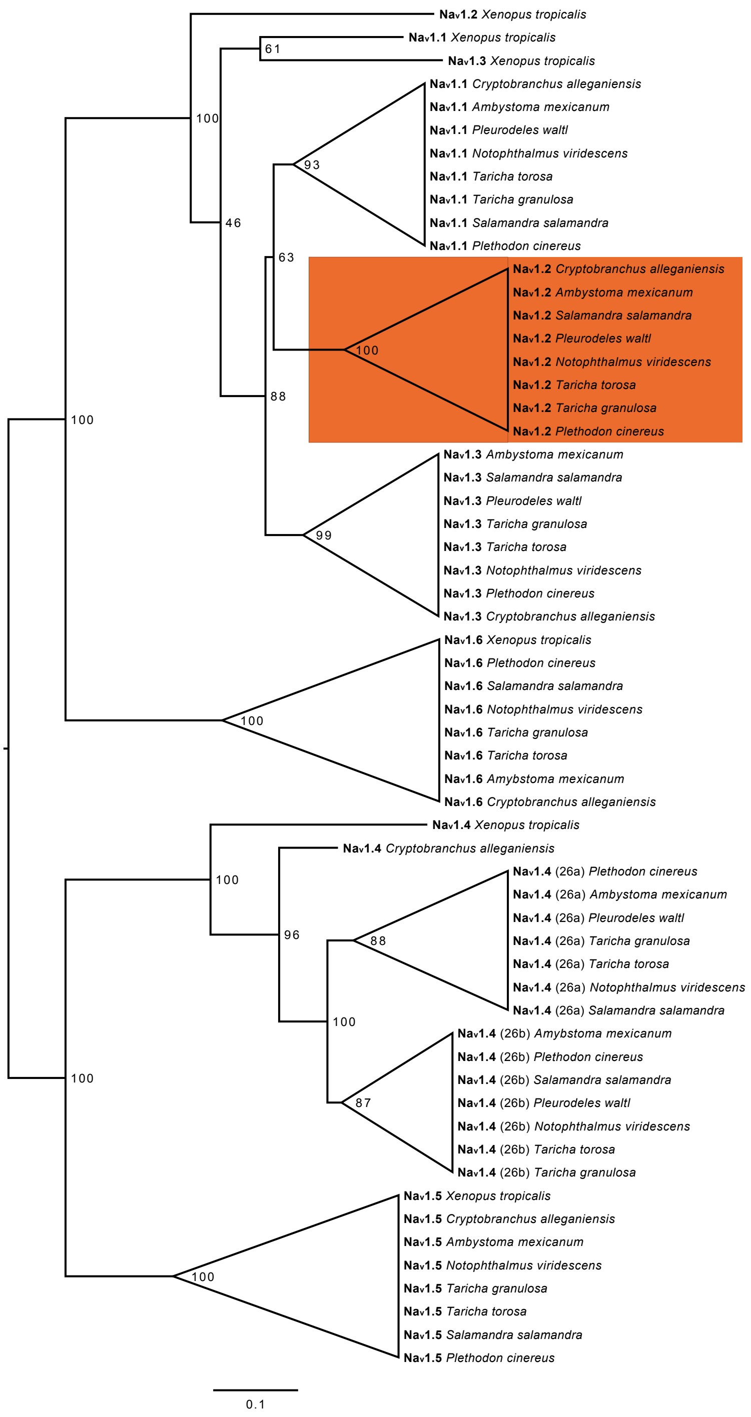


**Fig. S3.** Maximum likelihood tree constructed from 1063 bp nucleotide alignment of *SCNA* exon 26 sequences. Node labels indicate bootstrap support from 100 replicates. Orange highlighting indicates the clade grouping *SCN2A* from *Ambystoma* with *SCN2A* from other salamander species (bootstrap support 100%).

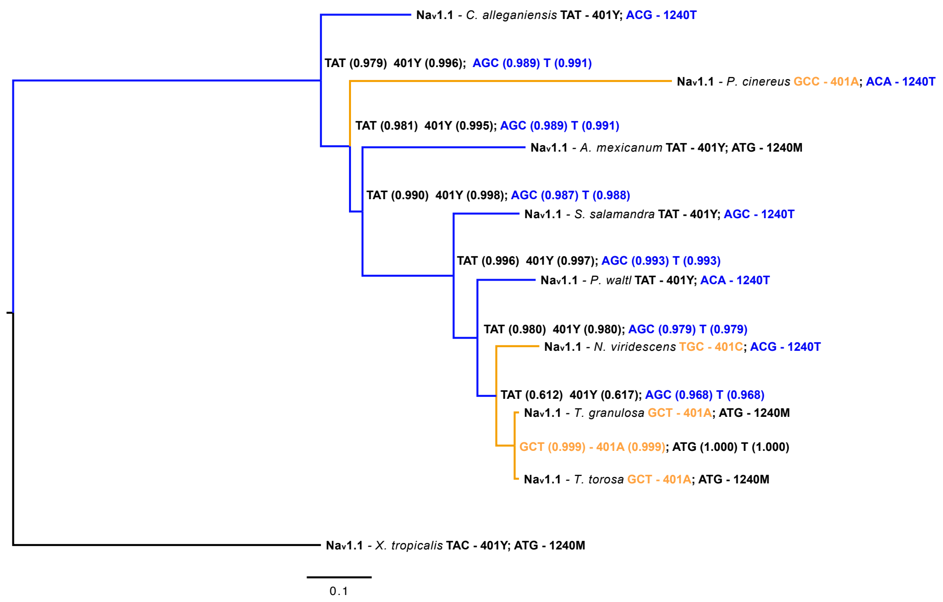


**Fig. S4.** Ancestral sequence reconstruction of tetrodotoxin resistant substitutions in $Na_v1.1$ based on neutral site model (M8a). Numbers in parentheses indicate posterior probability support for ancestral sequence reconstruction at nodes. Branch lengths indicate number of synonymous substitutions per codon. Blue branches – moderately resistant substitution, orange branches – highly resistant substitution.

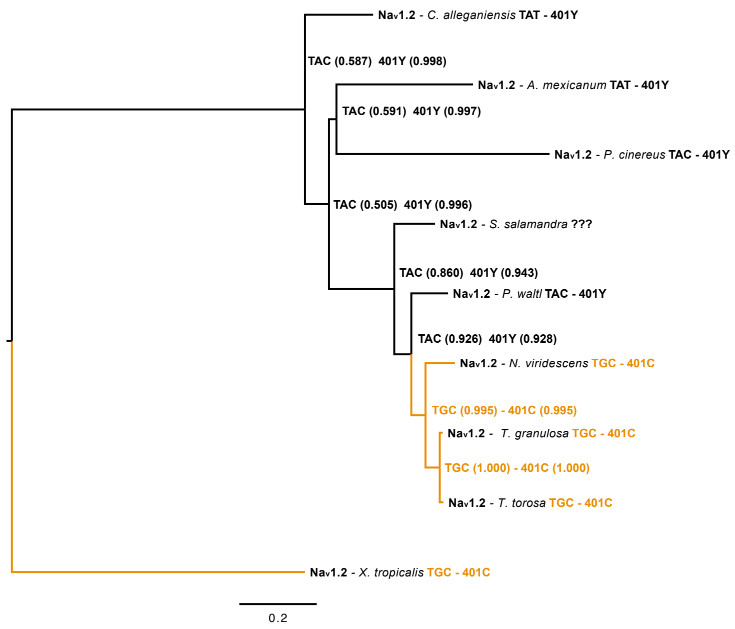


**Fig. S5.** Ancestral sequence reconstruction of tetrodotoxin resistant substitutions in $Na_v1.2$ based on neutral site model (M8a). Numbers in parentheses indicate posterior probability support for ancestral sequence reconstruction at nodes. Branch lengths indicate number of synonymous substitutions per codon. Blue branches – moderately resistant substitution, orange branches – highly resistant substitution.

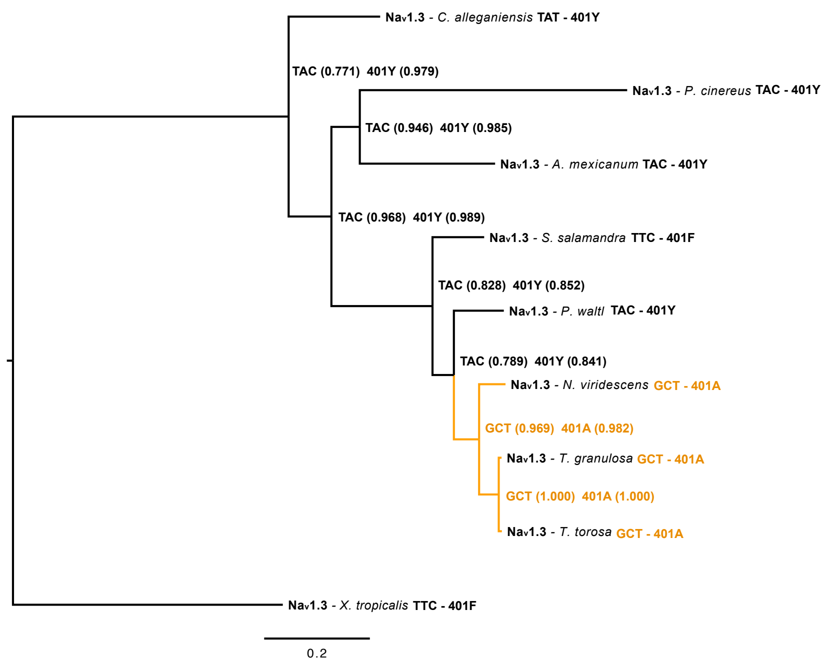


**Fig. S6.** Ancestral sequence reconstruction of tetrodotoxin resistant substitutions in $Na_v1.3$ based on site selection model (M8). Numbers in parentheses indicate posterior probability support for ancestral sequence reconstruction at nodes. Branch lengths indicate number of synonymous substitutions per codon. Blue branches – moderately resistant substitution, orange branches – highly resistant substitution.

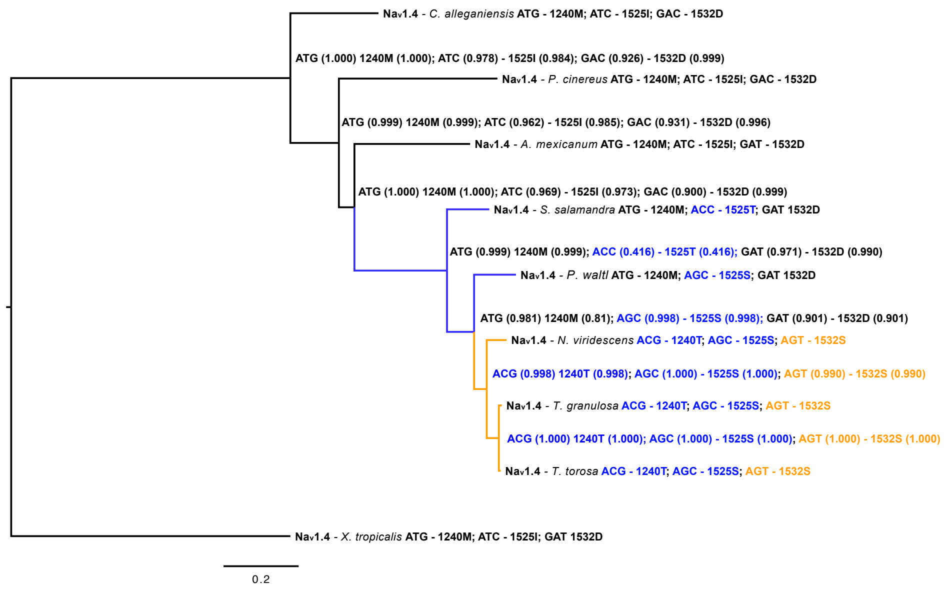


**Fig. S7.** Ancestral sequence reconstruction of tetrodotoxin resistant substitutions in $Na_v1.4$ based on neutral site model (M8a). Numbers in parentheses indicate posterior probability support for ancestral sequence reconstruction at nodes. Branch lengths indicate number of synonymous substitutions per codon. Blue branches – moderately resistant substitution, orange branches – highly resistant substitution.

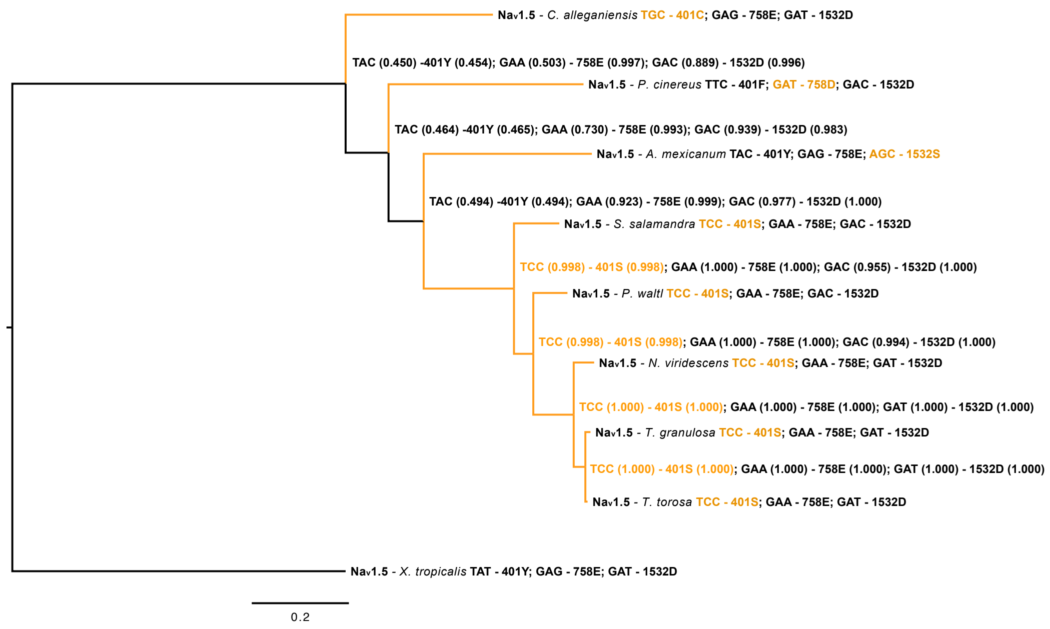


**Fig. S8.** Ancestral sequence reconstruction of tetrodotoxin resistant substitutions in $Na_v1.5$ based on neutral site model (M8a). Numbers in parentheses indicate posterior probability support for ancestral sequence reconstruction at nodes. Branch lengths indicate number of synonymous substitutions per codon. Blue branches – moderately resistant substitution, orange branches – highly resistant substitution.

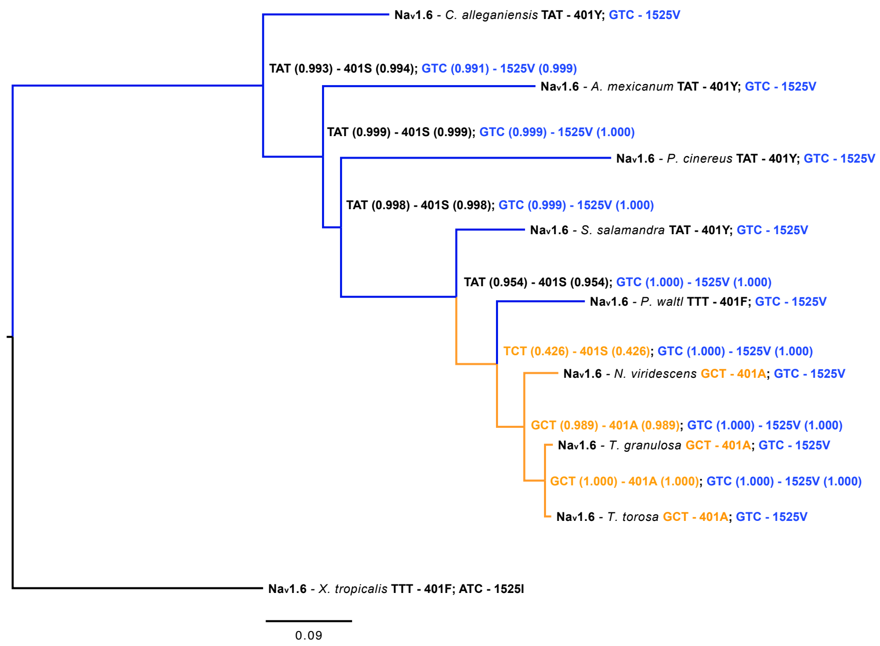


**Fig. S9.** Ancestral sequence reconstruction of tetrodotoxin resistant substitutions in $Na_v1.6$ based on neutral site model (M8a). Numbers in parentheses indicate posterior probability support for ancestral sequence reconstruction at nodes. Branch lengths indicate number of synonymous substitutions per codon. Blue branches – moderately resistant substitution, orange branches – highly resistant substitution.

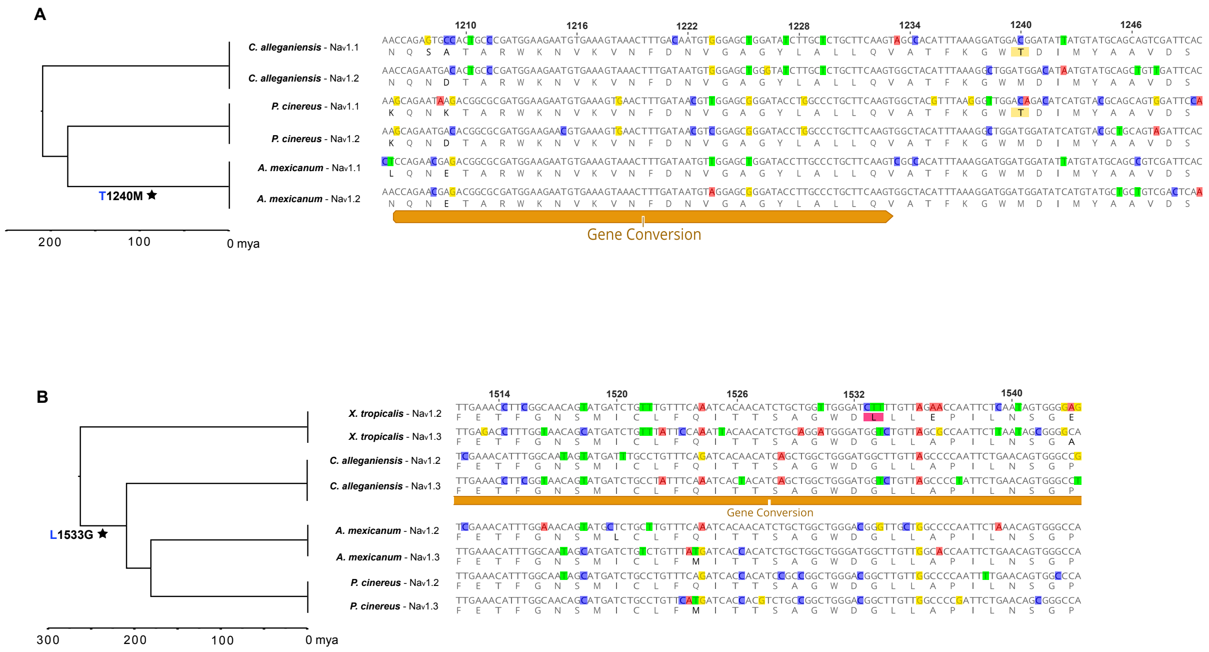


**Fig. S10.** Gene conversion within TTX binding regions associated with a loss of resistance-conferring substitutions. (A) Gene conversion was detected between the DIII P-loops of $Na_v1.1$ and $Na_v1.2$ in both *A. mexicanum*. The 1240T substitution conferring TTX resistance is highlighted in yellow. (B) Gene conversion was also detected between DIV P-loops of $Na_v1.2$ and $Na_v1.3$ within *C. alleganiensis*. The 1533L substitution putatively conferring TTX resistance is highlighted in pink. Site numbers are in reference to amino acids of rat $Na_v1.4$ (accession number AAA41682).

**Table S1.** Summary Statistics from Salamander $Na_v$ Sequencing and Alignments

| **SCN1A ($Na_v$1.1)** | **GC %** | **Length** | **Gaps** | **% Gaps** | **% Complete[a]** | **Pairwise % identity[b]** | **Average coverage** |
|---|---|---|---|---|---|---|---|
| *Ambystoma mexicanum* | 42.3 | 6224 | 165 | 2.7 | 97.3 | 81.8 | - |
| *Pleurodeles waltl* | 42.0 | 6149 | 390 | 6.3 | 93.7 | 81.9 | - |
| *Cryptobranchus alleganiensis* | 42.5 | 6216 | 162 | 2.6 | 97.4 | 81.6 | 21.44 |
| *Notophthalmus viridescens* | 41.5 | 6188 | 231 | 3.7 | 96.3 | 84.3 | 21.33 |
| *Taricha granulosa* | 41.5 | 6224 | 201 | 3.2 | 96.8 | 85.2 | 42.41 |
| *Taricha torosa* | 41.4 | 6224 | 159 | 2.6 | 97.4 | 85.4 | 39.88 |
| *Salamandra salamandra* | 41.6 | 5915 | 1846 | 31.2 | 68.8 | 60.2 | - |
| *Plethodon cinereus* | 46.1 | 6219 | 477 | 7.7 | 92.3 | 75.1 | 13.42 |
| *Xenopus tropicalis* | 41.2 | 6216 | 153 | 2.5 | 97.5 | 73.4 | - |
| ***SCN2A ($Na_v$1.2)*** | **GC %** | **Length** | **Gaps** | **% Gaps** | **% Complete** | **Pairwise % Identity** | **Average coverage** |
| *Ambystoma mexicanum* | 43.5 | 5946 | 402 | 6.8 | 93.2 | 70.9 | - |
| *Pleurodeles waltl* | 41.5 | 5982 | 108 | 1.8 | 98.2 | 79.7 | - |
| *Cryptobranchus alleganiensis* | 41.3 | 6000 | 438 | 7.3 | 92.7 | 74.9 | 18.6 |
| *Notophthalmus viridescens* | 42.3 | 6039 | 432 | 7.2 | 92.8 | 13.4 | 22.54 |
| *Taricha granulosa* | 41.8 | 6060 | 75 | 1.2 | 98.8 | 81.7 | 40.43 |
| *Taricha torosa* | 41.9 | 6060 | 69 | 1.1 | 98.9 | 81.6 | 34.32 |
| *Salamandra salamandra* | 41.9 | 3672 | 1077 | 29.3 | 70.7 | 56.8 | - |
| *Plethodon cinereus* | 46.7 | 6006 | 700 | 10 | 90.0 | 68.4 | 22.7 |
| *Xenopus tropicalis* | 41.4 | 6054 | 141 | 2.3 | 97.7 | 66.4 | - |
| ***SCN3A ($Na_v$1.3)*** | **GC %** | **Length** | **Gaps** | **% Gaps** | **% Complete** | **Pairwise % Identity** | **Average coverage** |
| *Ambystoma mexicanum* | 45 | 6072 | 51 | 0.8 | 99.2 | 83.3 | - |
| *Pleurodeles waltl* | 42.3 | 6069 | 195 | 3.2 | 96.8 | 85.9 | - |
| *Cryptobranchus alleganiensis* | 43.3 | 6093 | 39 | 0.6 | 99.4 | 83.7 | 23.59 |
| *Notophthalmus viridescens* | 42.4 | 6078 | 72 | 1.2 | 98.8 | 88.3 | 49.48 |
| *Taricha granulosa* | 42.3 | 6093 | 72 | 1.2 | 98.8 | 88.6 | 42.71 |
| *Taricha torosa* | 42.2 | 6099 | 72 | 1.2 | 98.8 | 88.6 | 40.9 |
| *Salamandra salamandra* | 42.9 | 6096 | 186 | 3.1 | 96.9 | 85.3 | - |
| *Plethodon cinereus* | 47.9 | 6096 | 510 | 8.4 | 91.6 | 75.1 | 14.44 |
| *Xenopus tropicalis* | 42.2 | 6093 | 36 | 0.6 | 99.4 | 75.5 | - |
| ***SCN4A ($Na_v$1.4)*** | **GC %** | **Length** | **Gaps** | **% Gaps** | **% Complete** | **Pairwise % Identity** | **Average coverage** |
| *Ambystoma mexicanum* | 43.4 | 5034 | 141 | 2.8 | 97.2 | 81.8 | - |
| *Pleurodeles waltl* | 43.3 | 5622 | 96 | 1.7 | 98.3 | 85.3 | - |
| *Cryptobranchus alleganiensis* | 46.2 | 5580 | 48 | 0.9 | 99.1 | 80.9 | 12.46 |
| *Notophthalmus viridescens* | 43.6 | 5622 | 96 | 1.7 | 98.3 | 87.1 | 47.29 |
| *Taricha granulosa* | 44 | 5469 | 87 | 1.6 | 98.4 | 87.3 | 35.46 |
| *Taricha torosa* | 44 | 5469 | 90 | 1.6 | 98.4 | 87.4 | 42.97 |
| *Salamandra salamandra* | 44.5 | 5619 | 135 | 2.4 | 97.6 | 85.4 | - |
| *Plethodon cinereus* | 43.5 | 5628 | 279 | 5 | 95 | 78.2 | 12.2 |
| *Xenopus tropicalis* | 43.3 | 5622 | 111 | 2 | 98 | 71.2 | - |
| ***SCN5A ($Na_v$1.5)*** | **GC %** | **Length** | **Gaps** | **% Gaps** | **% Complete** | **Pairwise % Identity** | **Average coverage** |
| *Ambystoma mexicanum* | 48.9 | 5988 | 45 | 0.8 | 99.2 | 82.7 | - |
| *Pleurodeles waltl* | 44.7 | 5706 | 102 | 1.8 | 98.2 | 86.9 | - |
| *Cryptobranchus alleganiensis* | 47 | 5961 | 33 | 0.6 | 99.4 | 81.6 | 16.06 |
| *Notophthalmus viridescens* | 45.5 | 5943 | 42 | 0.7 | 99.3 | 88.6 | 55.83 |
| *Taricha granulosa* | 45.7 | 5967 | 42 | 0.7 | 99.3 | 88.8 | 58.51 |
| *Taricha torosa* | 45.7 | 5967 | 42 | 0.7 | 99.3 | 89.0 | 36.8 |
| *Salamandra salamandra* | 45.7 | 5097 | 237 | 4.6 | 95.4 | 84.1 | - |
| *Plethodon cinereus* | 48.7 | 5961 | 48 | 0.8 | 99.2 | 81.8 | 22.49 |
| *Xenopus tropicalis* | 47 | 5988 | 45 | 0.8 | 99.2 | 73.3 | - |

| *SCN8A ($Na_v1.6$)* | GC % | Length | Gaps | % Gaps | % Complete | Pairwise % Identity | Average coverage |
|---|---|---|---|---|---|---|---|
| *Ambystoma mexicanum* | 44.5 | 5946 | 30 | 0.5 | 99.5 | 88.0 | - |
| *Pleurodeles waltl* | 44.5 | 5691 | 84 | 1.5 | 98.5 | 90.0 | - |
| *Cryptobranchus alleganiensis* | 44.7 | 5946 | 21 | 0.4 | 99.6 | 88.2 | 21.26 |
| *Notophthalmus viridescens* | 44.7 | 5916 | 30 | 0.5 | 99.5 | 91.5 | 49.65 |
| *Taricha granulosa* | 44.8 | 5952 | 30 | 0.5 | 99.5 | 91.7 | 43.78 |
| *Taricha torosa* | 44.8 | 5907 | 30 | 0.5 | 99.5 | 91.8 | 35.75 |
| *Salamandra salamandra* | 44.9 | 5946 | 30 | 0.5 | 99.5 | 90.8 | - |
| *Plethodon cinereus* | 44.7 | 5949 | 42 | 0.7 | 99.3 | 86.8 | 32.32 |
| *Xenopus tropicalis* | 41.9 | 5946 | 18 | 0.3 | 99.7 | 79.1 | - |

[a] Number of base pairs in a sequence divided by the total alignment length
[b] Average of all pairwise comparisons within an alignment

**Table S2.** Synonymous and Nonsynonymous Polymorphism in Salamander $Na_v$ Genes Sequenced for this Study

| | *SCN1A* | *SCN2A* | *SCN3A* | *SCN4A* | *SCN5A* | *SCN8A* |
|---|---|---|---|---|---|---|
| ***C. alleganiensis* (n=2)** | | | | | | |
| Synonymous ($P_s$) | 3 | 4 | 1 | 1 | 2 | 0 |
| Nonsynonymous ($P_n$) | 0 | 2 | 0 | 0 | 0 | 0 |
| Insertions | 0 | 0 | 0 | 0 | 0 | 0 |
| Missing exons | 0 | 1 | 0 | 0 | 0 | 0 |
| ***P. cinereus* (n=2)** | | | | | | |
| Synonymous ($P_s$) | 0 | 3 | 3 | 6 | 4 | 0 |
| Nonsynonymous ($P_n$) | 1 | 0 | 0 | 1 | 0 | 0 |
| Insertions | 0 | 0 | 0 | 0 | 0 | 0 |
| Missing exons | 0 | 2 | 2 | 1 | 0 | 0 |
| ***N. viridescens* (n=3)** | | | | | | |
| Synonymous ($P_s$) | 24 | 22 | 35 | 19 | 11 | 12 |
| Nonsynonymous ($P_n$) | 5 | 2 | 4 | 8 | 1 | 1 |
| Insertions | 1 (39 bases) | 0 | 0 | 0 | 0 | 0 |
| Missing exons | 0 | 2 | 0 | 0 | 0 | 0 |
| ***T. torosa* (n=3)** | | | | | | |
| Synonymous ($P_s$) | 2 | 5 | 2 | 2 | 1 | 0 |
| Nonsynonymous ($P_n$) | 1 | 2 | 0 | 1 | 1 | 0 |
| Insertions | 0 | 0 | 0 | 1 (3 bases) | 0 | 0 |
| Missing exons | 0 | 2 | 0 | 0 | 0 | 0 |
| ***T. granulosa* (n=3)** | | | | | | |
| Synonymous ($P_s$) | 4 | 8 | 3 | 10 | 4 | 5 |
| Nonsynonymous ($P_n$) | 0 | 0 | 0 | 2 | 2 | 0 |
| Insertions | 0 | 0 | 0 | 0 | 0 | 0 |
| Missing exons | 0 | 2 | 0 | 0 | 0 | 0 |

**Table S3.** Likelihood Ratio Tests from Codeml Site Models

| | $\ell$ | n Parameters | Parameter Estimates | $2\Delta\ell$ |
|---|---|---|---|---|
| ***SCN1A* ($\mathbf{Na_v1.1}$)** | | | | |
| Nearly neutral site model (M1a) | -20237.40 | 18 | $p_0$ = 0.91, ($p_1$ = 0.09) | - |
| | | | $\omega_0$ = 0.03, ($\omega_1$ = 1.00) | |
| Site selection model (M2a) | -20237.40 | 20 | $p_0$ = 0.91, $p_1$ = 0.09, ($p_2$ = 0.00) | (M2a vs. M1a) **0.00** |
| | | | $\omega_0$ = 0.03, ($\omega_1$ = 1.00), $\omega_2$ = 50.8 | |
| Site null model (M7) | -20200.36 | 18 | p = 0.11, q = 1.01 | - |
| Site neutral model (M8a) | -20193.38 | 19 | p = 0.13, q = 1.69 | - |
| | | | $p_0$ = 0.96, ($p_1$ = 0.04) | |
| Site selection model (M8) | -20193.38 | 20 | p = 0.13, q = 1.69 | (M7 vs. M8) **13.97**** |
| | | | $p_0$ = 0.96, ($p_1$ = 0.04), $\omega_1$ = 1.00 | (M8 vs. M8a) **0.00** |
| ***SCN2A* ($\mathbf{Na_v1.2}$)** | | | | |
| Nearly neutral site model (M1a) | -21559.11 | 18 | $p_0$ = 0.89, ($p_1$ = 0.11) | - |
| | | | $\omega_0$ = 0.05, ($\omega_1$ = 1.00) | |
| Site selection model (M2a) | -21559.11 | 20 | $p_0$ = 0.89, $p_1$ = 0.11, ($p_2$ = 0.00) | (M2a vs. M1a) **0.00** |
| | | | $\omega_0$ = 0.05, ($\omega_1$ = 1.00), $\omega_2$ = 45.2 | |
| Site null model (M7) | -21481.17 | 18 | p = 0.19, q = 1.43 | - |
| Site neutral model (M8a) | -21474.94 | 19 | p = 0.22, q = 2.14 | - |
| | | | $p_0$ = 0.97, ($p_1$ = 0.03) | |
| Site selection model (M8) | -21474.89 | 20 | p = 0.21, q = 2.07 | (M7 vs. M8) **12.56**** |
| | | | $p_0$ = 0.98, ($p_1$ = 0.02), $\omega_1$ = 1.09 | (M8 vs. M8a) **0.10** |
| ***SCN3A* ($\mathbf{Na_v1.3}$)** | | | | |
| Nearly neutral site model (M1a) | -21487.41 | 18 | $p_0$ = 0.91, ($p_1$ = 0.09) | - |
| | | | $\omega_0$ = 0.03, ($\omega_1$ = 1.00) | |
| Site selection model (M2a) | -21487.41 | 20 | $p_0$ = 0.91, $p_1$ = 0.09, ($p_2$ = 0.00) | (M2a vs. M1a) **2.92** |
| | | | $\omega_0$ = 0.03, ($\omega_1$ = 1.00), $\omega_2$ = 4.73 | |
| Site null model (M7) | -21439.44 | 18 | p = 0.12, q = 0.99 | - |
| Site neutral model (M8a) | -21406.75 | 19 | p = 0.14, q = 1.84 | - |
| | | | $p_0$ = 0.96, ($p_1$ = 0.04) | |
| Site selection model (M8) | -21401.10 | 20 | p = 0.13, q = 1.48 | (M7 vs. M8) **76.68**** |
| | | | $p_0$ = 0.98, ($p_1$ = 0.02), $\omega_1$ = 1.83 | (M8 vs. M8a) **11.30**** |
| ***SCN4A* ($\mathbf{Na_v1.4}$)** | | | | |
| Nearly neutral site model (M1a) | -21591.93 | 18 | $p_0$ = 0.83, ($p_1$ = 0.17) | - |
| | | | $\omega_0$ = 0.03, ($\omega_1$ = 1.00) | |
| Site selection model (M2a) | -21591.93 | 20 | $p_0$ = 0.83, $p_1$ = 0.17, ($p_2$ = 0.00) | (M2a vs. M1a) **0.00** |
| | | | $\omega_0$ = 0.03 ($\omega_1$ = 1.00), $\omega_2$ = 107.5 | |
| Site null model (M7) | -21528.57 | 18 | p = 0.14, q = 0.78 | - |
| Site neutral model (M8a) | -21521.16 | 19 | p = 0.16, q = 1.54 | - |
| | | | $p_0$ = 0.93, ($p_1$ = 0.07) | |
| Site selection model (M8) | -21521.09 | 20 | p = 0.16, q = 1.27 | (M7 vs. M8) **14.82**** |
| | | | $p_0$ = 0.94, ($p_1$ = 0.06), $\omega_1$ = 1.09 | (M8 vs. M8a) **0.14** |
| ***SCN5A* ($\mathbf{Na_v1.5}$)** | | | | |
| Nearly neutral site model (M1a) | -22525.65 | 18 | $p_0$ = 0.88 ($p_1$ = 0.12) | - |
| | | | $\omega_0$ = 0.04, ($\omega_1$ = 1.00) | |
| Site selection model (M2a) | -22523.34 | 20 | $p_0$ = 0.88, $p_1$ = 0.12, ($p_2$ = 0.00) | (M2a vs. M1a) **4.63** |
| | | | $\omega_0$ = 0.03 ($\omega_1$ = 1.00), $\omega_2$ = 30.1 | |
| Site null model (M7) | -22482.16 | 18 | p = 0.15, q = 1.04 | - |
| Site neutral model (M8a) | -22465.19 | 19 | p = 0.20, q = 2.28 | - |
| | | | $p_0$ = 0.95, ($p_1$ = 0.05) | |
| Site selection model (M8) | -22465.19 | 20 | p = 0.20, q = 2.28 | (M7 vs. M8) **33.94**** |
| | | | $p_0$ = 0.95, ($p_1$ = 0.05), $\omega_1$ = 1.00 | (M8 vs. M8a) **0.00** |

| | | | | |
|---|---|---|---|---|
| ***SCN8A* (Na$_v$1.6)** | | | | |
| Nearly neutral site model (M1a) | -18903.82 | 18 | $p_0$ = 0.94 ($p_1$ = 0.06)<br>$\omega_0$ = 0.02, ($\omega_1$ = 1.00) | - |
| Site selection model (M2a) | -18903.82 | 20 | $p_0$ = 0.94, $p_1$ = 0.06, ($p_2$ = 0.00)<br>$\omega_0$ = 0.02 ($\omega_1$ = 1.00), $\omega_2$ = 89.1 | (M2a vs. M1a) **0.00** |
| Site null model (M7) | -18911.33 | 18 | p = 0.09, q = 0.99 | - |
| Site neutral model (M8a) | -18879.57 | 19 | p = 0.12, q = 2.25<br>$p_0$ = 0.97, ($p_1$ = 0.03) | - |
| Site selection model (M8) | -18878.56 | 20 | p = 0.11, q = 1.70<br>$p_0$ = 0.98, ($p_1$ = 0.02), $\omega_1$ = 1.42 | (M7 vs. M8) **65.54****<br>(M8 vs. M8a) **2.02** |

** P-value < 0.01, determined by likelihood ratio test using $\chi^2$ distribution

* P-value < 0.05, determined by likelihood ratio test using $\chi^2$ distribution

**Table S4.** Likelihood Ratio Tests from Codeml Branch and Branch-Site Models Comparing Newts (Foreground) with Other Salamanders (Background)

| | $\ell$ | n Parameters | Parameter Estimates | $2\Delta\ell$ |
|---|---|---|---|---|
| ***SCN1A* (Na$_v$1.1)** | | | | |
| One ratio model (M0) | -20546.55 | 17 | $\omega$ = 0.07 | **-** |
| Branch model | -20526.69 | 18 | $\omega$ salamanders = 0.07, $\omega$ newts = 0.17 | (Branch vs. M0) **39.71**** |
| Branch-site neutral model (A1) | -20229.55 | 19 | $p_0$ = 0.86, $p_1$ = 0.08, ($p_2$ = 0.05) $\omega_0$ = 0.03, ($\omega_1$ = 1.00) | - |
| Branch-site selection model (A) | -20229.55 | 20 | $p_0$ = 0.86, $p_1$ = 0.08, ($p_2$ = 0.05) $\omega_0$ = 0.03, ($\omega_1$ = 1.00), $\omega_2$ = 1.00 | (A vs. A1) **0.00** |
| ***SCN2A* (Na$_v$1.2)** | | | | |
| One ratio model (M0) | -21838.97 | 17 | $\omega$ = 0.09 | **-** |
| Branch model | -21838.97 | 18 | $\omega$ salamanders = 0.09, $\omega$ newts = 0.09 | (Branch vs. M0) **0.00** |
| Branch-site neutral model (A1) | -21558.98 | 19 | $p_0$ = 0.89, $p_1$ = 0.11, ($p_2$ = 0.00) $\omega_0$ = 0.05, ($\omega_1$ = 1.00) | - |
| Branch-site selection model (A) | -21558.52 | 20 | $p_0$ = 0.89, $p_1$ = 0.11, ($p_2$ = 0.00) $\omega_0$ = 0.05, ($\omega_1$ = 1.00), $\omega_2$ = 7.84 | (A vs. A1) **0.94** |
| ***SCN3A* (Na$_v$1.3)** | | | | |
| One ratio model (M0) | -21867.12 | 17 | $\omega$ = 0.08 | **-** |
| Branch model | -21853.31 | 18 | $\omega$ salamanders = 0.07, $\omega$ newts = 0.15 | (Branch vs. M0) **27.63**** |
| Branch-site neutral model (A1) | -21471.92 | 19 | $p_0$ = 0.89, $p_1$ = 0.08, ($p_2$ = 0.03) $\omega_0$ = 0.03, ($\omega_1$ = 1.00) | - |
| Branch-site selection model (A) | -21468.13 | 20 | $p_0$ = 0.90, $p_1$ = 0.09, ($p_2$ = 0.01) $\omega_0$ = 0.03, ($\omega_1$ = 1.00), $\omega_2$ = 5.50 | (A vs. A1) **7.59*** |
| ***SCN4A* (Na$_v$1.4)** | | | | |
| One ratio model (M0) | -22097.03 | 17 | $\omega$ = 0.11 | **-** |
| Branch model | -22082.43 | 18 | $\omega$ salamanders = 0.10, $\omega$ newts = 0.23 | (Branch vs. M0) **29.20**** |
| Branch-site neutral model (A1) | -21576.31 | 19 | $p_0$ = 0.75, $p_1$ = 0.15, ($p_2$ = 0.10) $\omega_0$ = 0.03, ($\omega_1$ = 1.00) | - |
| Branch-site selection model (A) | -21576.16 | 20 | $p_0$ = 0.78, $p_1$ = 0.16, ($p_2$ = 0.06) $\omega_0$ = 0.03, ($\omega_1$ = 1.00), $\omega_2$ = 1.61 | (A vs. A1) **0.30** |
| ***SCN5A* (Na$_v$1.5)** | | | | |
| One ratio model (M0) | -22980.74 | 17 | $\omega$ = 0.09 | **-** |
| Branch model | -22978.73 | 18 | $\omega$ salamanders = 0.09, $\omega$ newts = 0.12 | (Branch vs. M0) **4.00** |
| Branch-site neutral model (A1) | -22525.65 | 19 | $p_0$ = 0.88, $p_1$ = 0.11, ($p_2$ = 0.00) $\omega_0$ = 0.02, ($\omega_1$ = 1.00) | - |
| Branch-site selection model (A) | -22525.65 | 20 | $p_0$ = 0.88, $p_1$ = 0.11, ($p_2$ = 0.00) $\omega_0$ = 0.02, ($\omega_1$ = 1.00), $\omega_2$ = 66.9 | (A vs. A1) **0.00** |
| ***SCN8A* (Na$_v$1.6)** | | | | |
| One ratio model (M0) | -19187.81 | 17 | $\omega$ = 0.06 | - |
| Branch model | -19183.65 | 18 | $\omega$ salamanders = 0.05, $\omega$ newts = 0.09 | (Branch vs. M0) **8.33*** |
| Branch-site neutral model (A1) | -18903.82 | 19 | $p_0$ = 0.94, $p_1$ = 0.06, ($p_2$ = 0.00) $\omega_0$ = 0.02, ($\omega_1$ = 1.00) | - |
| Branch-site selection model (A) | -18903.82 | 20 | $p_0$ = 0.94, $p_1$ = 0.06, ($p_2$ = 0.00) $\omega_0$ = 0.02, ($\omega_1$ = 1.00), $\omega_2$ = 1.00 | (A vs. A1) **0.00** |

** P-value < 0.01, determined by likelihood ratio test using $\chi^2$ distribution

* P-value < 0.05, determined by likelihood ratio test using $\chi^2$ distribution

**Table S5.** Posterior Probabilities for Sites with Elevated ω Values in Toxic Newts

| Site[a] | Exon | Nav1.1 | Nav1.2 | Nav1.3 | Nav1.4 | Nav1.5 | Nav1.6 |
|---|---|---|---|---|---|---|---|
| 12 | 1 | | | | 0.62 | | |
| 27 | 1 | | 0.91 | 0.55 | | | |
| 38 | 1 | | | | 0.52 | | |
| 120 | 2 | | | | | 0.77 | 0.75 |
| 155 | 3 | 0.53 | | | | | |
| 249 | 6 | 0.64 | | | | | |
| 278 | 6 | 0.58 | | | | | |
| 338 | 6 | | | 0.56 | | | |
| 340 | 6 | 0.79 | | | | | |
| 452 | 9 | | | 0.61 | | | |
| 460 | 9 | | | 0.60 | | | |
| 493 | 10 | 0.54 | | | | | |
| 543 | 13 | | | 0.98 | | | |
| 549 | 13 | | | 0.55 | | | |
| 621 | 14 | | | 0.59 | | | |
| 719 | 15 | | | 0.58 | 0.50 | | |
| **756[b]** | **15** | | | | **0.94** | | |
| **759[b]** | **15** | | | **0.61** | | | |
| 767 | 15 | | | 0.61 | | | |
| 774 | 15 | | | 0.66 | | | |
| 829 | 16 | 0.53 | | | | | |
| 837 | 16 | 0.60 | | | 0.54 | | |
| 842 | 16 | 0.59 | | | 0.51 | | |
| 843 | 16 | | | 0.51 | | | |
| 845 | 16 | 0.60 | | | | | |
| 877 | 16 | 0.51 | | | | | |
| 879 | 16 | | | 0.57 | | | |
| 881 | 16 | 0.60 | | | | | |
| 884 | 16 | | | 0.97 | | | |
| 887 | 16 | | | 0.60 | | | |
| 898 | 16 | | | 0.59 | | | |
| 911 | 16 | 0.61 | | | | | |
| 921 | 16 | 0.51 | | | | | 0.74 |
| 936 | 16 | 0.53 | | | | | |
| 940 | 16 | | | | | 0.69 | |
| 946 | 17 | 0.60 | | | | | |
| 957 | 17 | 0.58 | | | | | |
| 960 | 17 | 0.59 | | | | | |
| 965 | 17 | 0.95 | | | | | |
| 968 | 17 | 0.53 | | | | | |
| 981 | 17 | | | | 0.53 | | |
| 993 | 18 | | | | 0.52 | | |
| 1006 | 18 | | | 0.51 | 0.54 | | |
| 1028 | 18 | | | | 0.97 | | |
| 1046 | 19 | | | | 0.52 | | |
| 1127 | 20 | | | | | 0.56 | |
| 1179 | 21 | 0.59 | | | | | |
| 1187 | 21 | | | | 0.52 | | |
| 1189 | 21 | 0.51 | | | | | |
| 1191 | 21 | | | 0.86 | | | |
| 1194 | 21 | 0.60 | | 0.83 | | | |
| 1224 | 21 | 0.61 | | | | | |
| **1240[b]** | **22** | | | | **0.53** | | |
| 1250 | 22 | 0.61 | | | | | |
| 1254 | 23 | | | 0.54 | | | |
| 1257 | 23 | | | 0.97 | | | |
| 1261 | 23 | 0.52 | | | | | |
| 1262 | 23 | 0.61 | | | | | |
| 1367 | 25 | 0.60 | | | | | |
| 1383 | 25 | | | | 0.52 | | |
| 1390 | 25 | | | | 0.55 | | |
| **1519[b]** | **26** | | | | **0.52** | | |
| **1529[b]** | **26** | **0.52** | | | | | |
| **1532[b]** | **26** | | | | **0.98** | | |
| 1542 | 26 | | | 0.60 | | | |
| 1631 | 26 | 0.52 | | | | | |

| | | | | |
|---|---|---|---|---|
| 1737 | 26 | | 0.56 | |
| 1738 | 26 | | 0.51 | |
| 1739 | 26 | 0.53 | | |
| 1741 | 26 | | 0.51 | |
| 1744 | 26 | | 0.54 | |
| 1748 | 26 | | 0.52 | 0.53 |
| 1752 | 26 | | 0.83 | |
| 1774 | 26 | 0.61 | | |
| 1796 | 26 | 0.56 | | |
| 1817 | 26 | 0.59 | | |
| 1820 | 26 | 0.59 | | |
| 1827 | 26 | 0.51 | | |
| 1832 | 26 | 0.52 | | |
| 1939 | 26 | 0.52 | | |

[a] Site numbers are in reference to amino acid positions in the rat $Na_v1.4$ channel (accession number AAA41682)

[b] Known tetrodotoxin binding sites

**Table S6.** Posterior Probabilities for Sites Under Putative Positive Selection in All Salamanders

| Site[a] | Exon | $Na_v1.1$[b] | $Na_v1.2$[b] | $Na_v1.3$[b] | $Na_v1.4$[b] | $Na_v1.5$[b] | $Na_v1.6$[b] |
|---|---|---|---|---|---|---|---|
| 19 | 1 | | | 0.58; 0.81 | | | |
| 22 | 1 | | | | | | 0.70; 0.93 |
| 43 | 1 | | | 0.75; 0.95 | | | |
| 46 | 1 | | 0.51; 0.60 | | | | |
| 56 | 1 | | | 0.56; 0.79 | | | |
| 73 | 1 | 0.52; 0.63 | | 0.51 | | 0.60; 0.84 | 0.68; 0.91 |
| 74 | 1 | | | | | | 0.76; 0.96 |
| 80 | 1 | | | | | 0.54 | |
| 115 | 2 | | | | 0.93 | | |
| 155 | 3 | | | | | 0.52 | |
| 185 | 4 | | | | | 0.90; 0.99 | |
| 202 | 4 | | | 0.56 | | | |
| 209 | 5 | | | | | 0.62 | |
| 287 | 6 | 0.65; 0.88 | 0.51; 0.83 | | | | |
| 289 | 6 | 0.54 | | | | | |
| 290 | 6 | | 0.56; 0.67 | | | | |
| 292 | 6 | | | | 0.86 | | |
| 294 | 6 | 0.62; 0.85 | | | | 0.52 | |
| 295 | 6 | | | | | 0.67; 0.93 | |
| 297 | 6 | | | | 0.82 | | |
| 298 | 6 | 0.56; 0.78 | | | | 0.51; 0.71 | 0.60 |
| 300 | 6 | 0.53; 0.70 | 0.60; 0.82 | | | | |
| 301 | 6 | | | | | 0.56 | |
| 302 | 6 | 0.51; 0.62 | 0.56; 0.68 | 0.65; 0.88 | | | |
| 306 | 6 | | 0.71; 0.91 | | 0.67 | | |
| 307 | 6 | | | 0.68; 0.92 | | 0.53 | |
| 309 | 6 | | | 0.59; 0.81 | | 0.60 | |
| 311 | 6 | | | | 0.56 | 0.58 | |
| 325 | 6 | | | | 0.92 | | |
| 326 | 6 | | | | 0.77 | | |
| 328 | 6 | | 0.53 | | | | |
| 329 | 6 | | | 0.51; 0.66 | | | |
| 330 | 6 | | | | | 0.53 | |
| 332 | 6 | | | | | | 0.54; 0.72 |
| 333 | 6 | 0.66 | | 0.64; 0.88 | 0.63 | | 0.56; 0.77 |
| 337 | 6 | | 0.70; 0.90 | 0.81; 0.98 | | | |
| 338 | 6 | 0.55; 0.75 | | | | | |
| 339 | 6 | 0.61; 0.82 | 0.61; 0.75 | | | | 0.57; 0.79 |
| 340 | 6 | | | 0.62 | | 0.63 | 0.52; 0.61 |
| 344 | 7 | 0.60; 0.80 | 0.64; 0.82 | 0.51; 0.67 | | 0.55 | |
| 345 | 7 | | | | | 0.69; 0.93 | 0.52; 0.61 |
| 346 | 7 | | | | | 0.69; 0.94 | |
| 348 | 7 | | | 0.70; 0.92 | | 0.66; 0.90 | |
| 351 | 7 | 0.60; 0.81 | | | | | |
| 358 | 7 | 0.55; 0.71 | 0.58; 0.72 | | | | |
| 365 | 8 | | | 0.59; 0.80 | | 0.62; 0.88 | |
| 366 | 8 | | | | | 0.52; 0.73 | |
| 368 | 8 | 0.52; 0.70 | 0.56; 0.71 | | 0.63 | | 0.60; 0.82 |
| 374 | 8 | | | | | | 0.62 |
| **401**[c] | **8** | **0.61; 0.85** | | **0.51; 0.65** | | | **0.51; 0.68** |
| 423 | 9 | | | | 0.51 | | |
| 476 | 10 | | | | 0.57 | | |
| 485 | 10 | 0.51; 0.64 | | | 0.76 | | |
| 486 | 10 | | | | 0.61 | | |
| 487 | 10 | 0.53; 0.71 | | | | | |
| 492 | 10 | | | | 0.65 | | |
| 505 | 10 | | | | 0.55 | | |
| 521 | 10 | | | | 0.79 | | |
| 555 | 13 | 0.58; 0.74 | 0.53; 0.61 | 0.78; 0.97 | | | |
| 557 | 13 | 0.60 | | 0.68; 0.91 | 0.87 | | 0.53; 0.62 |
| 560 | 13 | 0.65 | | | | | |
| 563 | 13 | 0.66; 0.88 | | | | 0.56 | |
| 567 | 13 | | | | | | 0.51 |
| 598 | 13 | 0.62; 0.87 | | | 0.56 | 0.73; 0.96 | 0.57; 0.81 |
| 601 | 13 | | | 0.80; 0.97 | | | |
| 602 | 13 | | | | | 0.66; 0.91 | 0.65; 0.85 |

| | | | | | | | |
|---|---|---|---|---|---|---|---|
| 606 | 13 | | 0.52 | | | | |
| 609 | 13 | | | | | 0.53 | |
| 654 | 14 | | | 0.76; 0.96 | | | |
| 728 | 15 | | 0.57; 0.65 | | | | |
| 729 | 15 | | | 0.51; 0.64 | 0.97 | 0.50; 0.71 | 0.75; 0.96 |
| 732 | 15 | | 0.52; 0.56 | 0.61; 0.81 | 0.78 | 0.71; 0.94 | |
| 739 | 15 | | | | | 0.54; 0.79 | |
| 774 | 15 | | 0.70; 0.90 | | | | |
| 828 | 16 | | | | | 0.70 | |
| 830 | 16 | | | | | | 0.87; 0.99 |
| 832 | 16 | 0.55 | 0.57 | | | | 0.85; 0.99 |
| 840 | 16 | | | | | | 0.53; 0.71 |
| 841 | 16 | | | | 0.73 | 0.64 | |
| 846 | 16 | | | | | 0.61 | |
| 848 | 16 | | | | | 0.57; 0.82 | |
| 849 | 16 | | | | | | 0.51 |
| 850 | 16 | | | | | 0.63 | |
| 852 | 16 | | | 0.56; 0.69 | | 0.55; 0.80 | |
| 864 | 16 | | | 0.54; 0.66 | | | |
| 877 | 16 | | | | | 0.68; 0.93 | |
| 878 | 16 | | | | | 0.61 | |
| 881 | 16 | | | | 0.81 | 0.51 | |
| 886 | 16 | | | | | 0.52; 0.76 | |
| 887 | 16 | | 0.58 | | | | |
| 899 | 16 | | | | | 0.54 | |
| 912 | 16 | | 0.60; 0.82 | | | | |
| 913 | 16 | | 0.64; 0.82 | | | | |
| 916 | 16 | | | | 0.50 | | |
| 943 | 17 | | | | 0.55 | | |
| 950 | 17 | | | | | 0.64 | |
| 951 | 17 | | | 0.73; 0.94 | | | |
| 952 | 17 | 0.55 | | | | | |
| 967 | 17 | 0.53; 0.70 | | | | | |
| 971 | 17 | | | | 0.96 | | |
| 972 | 17 | 0.69; 0.91 | | | 0.67 | | |
| 976 | 17 | | | | | 0.53 | |
| 978 | 17 | 0.58; 0.79 | | | | | |
| 979 | 17 | | | 0.86; 0.99 | | | |
| 980 | 17 | | | | | 0.52; 0.72 | |
| 981 | 17 | 0.53; 0.71 | | | | | |
| 985 | 17 | | | | | | 0.64 |
| 997 | 18 | | | | 0.82 | | |
| 999 | 18 | 0.59 | | | | | |
| 1003 | 18 | | | | | 0.61 | |
| 1004 | 18 | | | 0.76; 0.96 | | | |
| 1006 | 18 | | | | | 0.68; 0.92 | |
| 1008 | 18 | | | | 0.61 | | |
| 1009 | 18 | | 0.64; 0.85 | | | 0.52; 0.74 | 0.77; 0.97 |
| 1012 | 18 | | | | | 0.52; 0.75 | |
| 1106 | 20 | | | | | 0.55; 0.79 | |
| 1111 | 20 | | | | | 0.64; 0.88 | |
| 1113 | 20 | | | | | 0.62; 0.85 | |
| 1115 | 20 | | | | | 0.65 | |
| 1187 | 21 | | | 0.50; 0.66 | | | |
| 1188 | 21 | | | | | 0.57; 0.82 | |
| 1189 | 21 | | | 0.54; 0.68 | | | 0.53 |
| 1192 | 21 | | | 0.72; 0.93 | | | |
| 1193 | 21 | 0.70; 0.93 | 0.63; 0.86 | 0.87; 0.99 | | | 0.73; 0.93 |
| 1194 | 21 | | | | 0.57 | | |
| 1195 | 21 | | | 0.57; 0.82 | | | |
| 1203 | 21 | 0.59; 0.78 | | | | | 0.64; 0.88 |
| 1204 | 21 | 0.58 | | 0.53 | | | 0.69; 0.92 |
| 1207 | 21 | 0.60; 0.81 | | 0.88; 0.99 | 0.76 | 0.61; 0.86 | |
| 1208 | 21 | | | 0.60; 0.81 | | | 0.55; 0.74 |
| 1211 | 21 | | | | 0.92 | | |
| 1216 | 21 | | | | | 0.58 | |
| 1251 | 22 | | | 0.67; 0.88 | | | |
| 1253 | 23 | 0.57; 0.80 | | | | | |
| 1254 | 23 | 0.59; 0.81 | | | | | |
| 1257 | 23 | | | | 0.89 | | |

| | | | | | | | |
|---|---|---|---|---|---|---|---|
| 1260 | 23 | | 0.52 | | | | |
| 1332 | 24 | 0.64; 0.85 | 0.53; 0.66 | | | | |
| 1334 | 25 | | 0.65; 0.87 | | | | |
| 1351 | 25 | | | | 0.67 | | |
| 1372 | 25 | | | | 0.72 | | 0.69; 0.92 |
| 1380 | 25 | 0.57 | | 0.69 | | | |
| 1390 | 25 | | | | | 0.62 | |
| 1424 | 25 | | | | 0.61 | | |
| **1533[c]** | **26** | **0.50; 0.67** | **0.55; 0.71** | | | | |
| 1542 | 26 | | | | | 0.66 | |
| 1543 | 26 | 0.67; 0.90 | | | | 0.70 | |
| 1549 | 26 | 0.50; 0.66 | | 0.66; 0.89 | | | |
| 1550 | 26 | | | | 0.89 | | |
| 1551 | 26 | | 0.58; 0.78 | 0.68; 0.91 | | 0.57; 0.80 | |
| 1553 | 26 | | | | 0.54 | | |
| 1556 | 26 | | | | 0.95 | | |
| 1558 | 26 | | | | 0.95 | 0.56 | |
| 1618 | 26 | | | | 0.54 | | |
| 1623 | 26 | | 0.56; 0.78 | | | | |
| 1631 | 26 | | | 0.53 | | | |
| 1635 | 26 | 0.53 | | | | | |
| 1725 | 26 | 0.60 | | | | 0.58; 0.83 | |
| 1726 | 26 | | 0.62; 0.81 | | | | |
| 1733 | 26 | 0.62; 0.81 | | | | | |
| 1736 | 26 | | | | 0.53 | | |
| 1739 | 26 | | 0.56; 0.70 | | | | |
| 1742 | 26 | | | | 0.66 | | |
| 1745 | 26 | | 0.53; 0.66 | | | | |
| 1746 | 26 | | 0.70; 0.92 | | 0.91 | | |
| 1747 | 26 | | 0.62; 0.83 | | 0.81 | | |
| 1748 | 26 | | 0.63; 0.84 | | | | |
| 1749 | 26 | | | | 0.71 | | |
| 1751 | 26 | | | | 0.63 | | |
| 1752 | 26 | | | | 0.61 | | |
| 1754 | 26 | | | | 0.93 | | |
| 1755 | 26 | | 0.62; 0.79 | 0.57 | | | |
| 1757 | 26 | | | | | | |
| 1760 | 26 | | | | 0.74 | | |
| 1767 | 26 | | | | 0.61 | | |
| 1769 | 26 | | | 0.58 | 0.61 | | |
| 1773 | 26 | | | | 0.57 | | |
| 1780 | 26 | | | | 0.78 | | |
| 1782 | 26 | | | | 0.60 | | |
| 1784 | 26 | | | | 0.74 | | |
| 1789 | 26 | | | | 0.87 | | |
| 1798 | 26 | | | | 0.69 | | |
| 1802 | 26 | | | | 0.87 | | |
| 1808 | 26 | | | | 0.77 | | |
| 1809 | 26 | | | | 0.51 | | |
| 1816 | 26 | | | | 0.67 | | |
| 1817 | 26 | | | | 0.51 | | |
| 1819 | 26 | | | | 0.69 | | |
| 1822 | 26 | | | 0.58; 0.82 | | | |
| 1828 | 26 | | | | | 0.62 | |
| 1835 | 26 | | | | 0.98 | | |
| 1839 | 26 | | | | 0.58 | | |

[a] Site numbers are in reference to amino acid positions in the rat $Na_v1.4$ channel (accession number AAA41682)

[b] Numbers indicate posterior probabilities of positive selection from empirical Bayes estimates in PAML. A single value indicates detection from the M8 model only and two values indicate detection from both the M2a model and M8 models.

[c] Known tetrodotoxin binding sites

**Table S7.** Amphibian $Na_v$ Sequences Used for Targeted NGS Probe Design

| Species | Source[a] | Best BLAST hit | Accession |
|---|---|---|---|
| *Ambystoma mexicanum* | WGS NCBI | *SCN1A* | gb\|PGSH01113157.1 |
| *Ambystoma mexicanum* | WGS NCBI | *SCN2A*, *SCN3A* | gb\|PGSH01109388.1 |
| *Ambystoma mexicanum* | WGS NCBI | *SCN4A* | gb\|PGSH01101866.1, gb\|PGSH01095590.1, gb\|JXRH01331098.1 |
| *Ambystoma mexicanum* | WGS NCBI | *SCN5A* | gb\|PGSH01008813.1 |
| *Ambystoma mexicanum* | WGS NCBI | *SCN8A* | gb\|PGSH01049067.1 |
| *Hynobius chinensis* | TSA NCBI | *SCN1A* | gb\|GAQK01012416.1, gb\|GAQK01089723.1, gb\|GAQK01123640.1, gb\|GAQK01022956.1, gb\|GAQK01037701.1, gb\|GAQK01110837.1, gb\|GAQK01035933.1, gb\|GAQK01049457.1, gb\|GAQK01026323.1 gb\|GAQK01118202.1, gb\|GAQK01062486.1, gb\|GAQK01026323.1, gb\|GAQK01062486.1 |
| *Hynobius chinensis* | TSA NCBI | *SCN2A* | gb\|GAQK01012415.1, gb\|GAQK01089724.1, gb\|GAQK01123639.1, gb\|GAQK01062305.1, gb\|GAQK01047585.1, gb\|GAQK01096581.1, gb\|GAQK01044980.1, gb\|GAQK01086119.1, gb\|GAQK01096592.1, gb\|GAQK01106518.1, gb\|GAQK01122588.1 |
| *Hynobius chinensis* | TSA NCBI | *SCN4A* | gb\|GAQK01140156.1, gb\|GAQK01024534.1, gb\|GAQK01021831.1, gb\|GAQK01021830.1, gb\|GAQK01082803.1, gb\|GAQK01071419.1, gb\|GAQK01071419.1 |
| *Hynobius chinensis* | TSA NCBI | *SCN5A* | gb\|GAQK01128205.1, gb\|GAQK01083790.1, gb\|GAQK01027263.1, gb\|GAQK01027805.1, gb\|GAQK01015614.1, gb\|GAQK01012146.1, gb\|GAQK01014539.1, gb\|GAQK01014539.1 |
| *Hynobius chinensis* | TSA NCBI | *SCN8A* | gb\|GAQK01067521.1, gb\|GAQK01020756.1, gb\|GAQK01038507.1, gb\|GAQK01045113.1, gb\|GAQK01022217.1, gb\|GAQK01096591.1, gb\|GAQK01071418.1, gb\|GAQK01071418.1 |
| *Hynobius retardatus* | TSA NCBI | *SCN1A* | gb\|LE210884.1, gb\|LE107081.1, gb\|LE105972.1 |
| *Hynobius retardatus* | TSA NCBI | *SCN3A* | gb\|LE175129.1 |
| *Hynobius retardatus* | TSA NCBI | *SCN4A* | gb\|LE175126.1, gb\|LE175128.1 |
| *Hynobius retardatus* | TSA NCBI | *SCN5A* | gb\|LE143587.1, gb\|LE143588.1 |
| *Hynobius retardatus* | TSA NCBI | *SCN8A* | gb\|LE175125.1 |

| | | | |
|---|---|---|---|
| *Lyciasalamandra atifi* | Transcriptome assembly provided by Miguel Vences (Rodríguez et al. 2017) | - | - |
| *Nanorana parkeri* | WGS NCBI | *SCN1A, SCN2A, SCN3A* | gb\|NW_017306417.1 |
| *Nanorana parkeri* | WGS NCBI | *SCN4A* | gb\|NW_017306748.1 |
| *Nanorana parkeri* | WGS NCBI | *SCN5A* | gb\|NW_017306389.1 |
| *Nanorana parkeri* | WGS NCBI | *SCN8A* | gb\|NW_017307114.1 |
| *Notophthalmus viridescens* | http://sandberg.cmb.ki.se/redspottednewt/ | - | - |
| *Paramesotriton hongkonginensis* | Sequence Read Archive NCBI SRX796492 | - | - |
| *Pleurodeles waltl* | Whole genome assembly provided by Ahmed Elewa (Elewa et al. 2017) | *SCN1A* | abyss_v4.2_66066951 |
| *Pleurodeles waltl* | Transcriptome assembly from iNewt Database: http://www.nibb.ac.jp/imori/main/ | *SCN1A* | TRINITY_DN288824_c1_g3_i7 |
| *Pleurodeles waltl* | Whole genome assembly provided by Ahmed Elewa (Elewa et al. 2017) | *SCN2A* | abyss_v4.2_66112789 |
| *Pleurodeles waltl* | Whole genome assembly provided by Ahmed Elewa (Elewa et al. 2017) | *SCN3A* | abyss_v4.2_66060341 |
| *Pleurodeles waltl* | Transcriptome assembly from iNewt Database: http://www.nibb.ac.jp/imori/main/ | *SCN3A* | TRINITY_DN288824_c1_g3_i6 |
| *Pleurodeles waltl* | Whole genome assembly provided by Ahmed Elewa (Elewa et al. 2017) | *SCN4A* | abyss_v4.2_48183054 |
| *Pleurodeles waltl* | Transcriptome assembly from iNewt Database: http://www.nibb.ac.jp/imori/main/ | *SCN4A* | TRINITY_DN288824_c1_g2_i5 |
| *Pleurodeles waltl* | Whole genome assembly provided by Ahmed Elewa (Elewa et al. 2017) | *SCN5A* | abyss_v4.2_66164693 |
| *Pleurodeles waltl* | Whole genome assembly provided by Ahmed Elewa (Elewa et al. 2017) | *SCN8A* | abyss_v4.2_66123907 |
| *Salamandra atra* | Transcriptome assembly provided by Miguel Vences (Rodríguez et al. 2017) | - | - |
| *Salamandra infraimmaculata* | Transcriptome assembly provided by Miguel Vences (Rodríguez et al. 2017) | - | - |
| *Salamandra salamandra* | TSA NCBI | *SCN1A* | gb\|GIKK01030996.1, gb\|GIKK01027377.1, gb\|GIKK01026170.1, gb\|GIKK01027688.1 |
| *Salamandra salamandra* | TSA NCBI | *SCN2A* | gb\|GIKK01012950.1, gb\|GIKK01007670.1, gb\|GIKK01006682.1 |
| *Salamandra salamandra* | TSA NCBI | *SCN3A* | gb\|GIKK01031859.1 |
| *Salamandra salamandra* | TSA NCBI | *SCN4A* | gb\|GIKK01007017.1, gb\|GIKK01015583.1 |
| *Salamandra salamandra* | TSA NCBI | *SCN5A* | gb\|GIKK01011042.1 |
| *Salamandra salamandra* | TSA NCBI | *SCN8A* | gb\|GIKK01019313.1, gb\|GIKK01023854.1, gb\|GIKK01002548.1 |
| *Tylototriton wenxianensis* | TSA NCBI | *SCN4A* | gb\|GESS01000732.1, gb\|GESS01024789.1, gb\|GESS01029581.1, gb\|GESS01063809.1 |

| | | | |
|---|---|---|---|
| *Tylototriton wenxianensis* | TSA NCBI | *SCN5A* | gb\|GESS01016882.1, gb\|GESS01035197.1 |
| *Xenopus tropicalis* | WGS NCBI | *SCN1A* | gb\|AAMC03035440.1 |
| *Xenopus tropicalis* | WGS NCBI | *SCN2A* | gb\|AAMC03035445.1 |
| *Xenopus tropicalis* | WGS NCBI | *SCN3A* | gb\|AAMC03035458.1, gb\|AAMC03035459.1, gb\|AAMC03035460.1 |
| *Xenopus tropicalis* | WGS NCBI | *SCN4A* | gb\|AAMC03036452.1 |
| *Xenopus tropicalis* | WGS NCBI | *SCN5A* | gb\|AAMC03022243.1, gb\|AAMC03022245.1, gb\|AAMC03022246.1, gb\|AAMC03022247.1, gb\|AAMC03022249.1, gb\|AAMC03022250.1, gb\|AAMC03022251.1 |
| *Xenopus tropicalis* | WGS NCBI | *SCN8A* | gb\|AAMC03008918.1, gb\|AAMC03008917.1, gb\|AAMC03008916.1 |

[a] WGS – whole genome shotgun database, TSA – transcriptome shotgun assembly database
